# The interaction between random and systematic visual stimulation and infraslow quasiperiodic spatiotemporal patterns of whole brain activity

**DOI:** 10.1101/2022.12.06.519337

**Authors:** Nan Xu, Derek M. Smith, George Jeno, Dolly T. Seeburger, Eric H. Schumacher, Shella D. Keilholz

**Affiliations:** Wallace H. Coulter Department of Biomedical Engineering, Georgia Institute of Technology and Emory University, Atlanta, GA, United States; School of Psychology, Georgia Institute of Technology, Atlanta, GA, United States; School of Computer Science, Georgia Institute of Technology, Atlanta, GA, United States; Department of Neurology, Division of Cognitive Neurology/Neuropsychology, The Johns Hopkins University School of Medicine, Baltimore, Maryland, United States

## Abstract

One prominent feature of the infraslow BOLD signal during rest or task is quasi-periodic spatiotemporal pattern (QPP) of signal changes that involves an alternation of activity in key functional networks and propagation of activity across brain areas, and that is known to tie to the infraslow neural activity involved in attention and arousal fluctuations. This ongoing whole-brain pattern of activity might potentially modify the response to incoming stimuli or be modified itself by the induced neural activity. To investigate this, we presented checkerboard sequences flashing at 6Hz to subjects. This is a salient visual stimulus that is known to produce a strong response in visual processing regions. Two different visual stimulation sequences were employed, a systematic stimulation sequence in which the visual stimulus appeared every 20.3 secs and a random stimulation sequence in which the visual stimulus occurred randomly every 14~62.3 secs. Three central observations emerged. First, the two different stimulation conditions affect the QPP waveform in different aspects, i.e., systematic stimulation has greater effects on its phase and random stimulation has greater effects on its magnitude. Second, the QPP was more frequent in the systematic condition with significantly shorter intervals between consecutive QPPs compared to the random condition. Third, the BOLD signal response to the visual stimulus across both conditions was swamped by the QPP at the stimulus onset. These results provide novel insights into the relationship between intrinsic patterns and stimulated brain activity.

## Introduction

Spontaneous fluctuations in blood oxygen level-dependent (BOLD) signals, recorded by functional magnetic resonance imaging (fMRI), capture the hemodynamic response to neural activity. The infraslow (less than 0.1Hz) BOLD fluctuations are suggested to have unique functional and neurophysiological principles that are distinct from higher frequencies (Chen et al., 2020; Grooms et al., 2017; Majeed et al., 2009, 2011; Mitra et al., 2018; Pan et al., 2013; Thompson et al., 2014). The spatiotemporal structure of the infraslow BOLD fluctuations has provided novel insights into the large-scale functional architecture of the brain, as well as its changes during tasks engagement, development, and disease (Fox & Raichle, 2007).

One type of spatiotemporal structure consists of a reproducible pattern of spatial changes that repeat over time, exhibiting an alternation of high and low activity in particular areas and propagation of activity along the cortex. These phase-locked quasi-periodic patterns (QPPs) are found to characterize the intrinsic dynamics of infraslow BOLD fluctuations in human brains (Bolt et al., 2022; Yousefi & Keilholz, 2021). The primary (or the strongest) QPP, in particular, displays prominent anticorrelation between the default mode network (DMN) and task-positive network (TPN) across rodents and humans (Belloy, Naeyaert, et al., 2018; Majeed et al., 2011; Raut et al., 2021; Yousefi et al., 2018; Yousefi & Keilholz, 2021). It has been shown to correlate with the infraslow neural activity (Grooms et al., 2017; Thompson et al., 2014), which is known to be involved in attention (Helps et al., 2010; Monto et al., 2008) and arousal (Raut et al., 2021; Sihn & Kim, 2022). The primary QPP can be affected by sustained attention and other attention control/working memory tasks (Abbas, Bassil, et al., 2019; Abbas, Belloy, et al., 2019), as well as arousal fluctuations (Raut et al., 2021).

The interactions between the infraslow activity and task- or stimulation-evoked brain responses have been the focus of much research over the last decade. Several studies (Chen et al., 2020; Fox et al., 2005; He, 2013; Huang et al., 2017) reveal that stimulation-evoked BOLD responses are affected by the magnitude of the spontaneous BOLD fluctuations at stimulus onsets, namely, *the prestimulus baseline*, which causes the widely observed *intra*-subject trial-to-trial variability in BOLD responses. However, revealed by Chen and his colleagues, the power of the evoked infraslow hemodynamics appeared to occur before the power of neural dynamics (Chen et al., 2020, Fig. 3ef), which implies that a significant portion of the hemodynamics may not directly arise from the neural level. Because the primary QPP captures the major dynamics of the intrinsic infraslow brain activity, by investigating stimulus-evoked QPPs, one may probe the interaction between the stimulation-evoked BOLD response and the spontaneous infraslow neural activity. A recent investigation in stimulation-evoked QPPs was demonstrated in anesthetized mice (Belloy et al., 2021), which suggests that visual stimulation can trigger the onsets of primary QPPs and that primary QPPs with different phases prior to the visual stimulus affect the magnitude of the subsequent visual response. Despite this progress, more remains to be investigated. Specifically, it is still unclear how environmental perturbations affect the ongoing QPPs and how ongoing QPPs modulate the BOLD responses to these environmental perturbations in humans.

Expanding upon previous findings, here we describe a comprehensive investigation into the relationship between the primary QPPs and visual stimulation in humans. Given that primary QPPs associated with both attention (Abbas, Bassil, et al., 2019; Abbas, Belloy, et al., 2019) and arousal fluctuations (Raut et al., 2021), we aimed to explore how QPPs interact with visual stimuli under conditions that held arousal fluctuations constant. We employed two conditions of different sequences of visual stimulation induced by flickering checkerboard flashing at 6Hz (which is unlikely to affect arousal levels), one involving a systematic stimulation sequence where the visual stimulus appeared every 20 seconds, and the other involving a random stimulation sequence where the visual stimulus occurred randomly between 14~62.3 seconds (average 19.95s+/− 6.37s). Notably, a systematic sequence has been routinely employed for evoking the infraslow spontaneous BOLD signals in previous studies (Belloy et al., 2021; Duann et al., 2002). These sequences have been shown to entrain the intrinsic rhythms of low-frequency brain oscillation to the structure of the attended stimulus stream (Ding et al., 2006; Lakatos et al., 2008) and enhance attention (Ding et al., 2006; Jones et al., 2002; Lakatos et al., 2008; Qiao et al., 2022). While our experiment only involved a basic sensory stimulation paradigm without requiring any responses or measuring attention behaviors, our findings may provide novel insights into how attentional processes are affected by sensory stimulation for future studies.

Given that QPPs are closely associated with infraslow neural activity (Grooms et al., 2017; Thompson et al., 2014), the intervals between predetermined stimuli were specially designed to ensure the frequencies of the presentation of these noninvasive stimuli within the infraslow range, i.e., 0.049Hz for the systematic and 0.016Hz~0.07Hz for random stimulation, that was suggested to modulate the infraslow neural fluctuations (Qiao et al., 2022). Additionally, an equal number of stimuli were presented across both systematic and random sequences. Hence, we can investigate the interaction between QPP and the stimuli and compare the results across two visual stimulation conditions. Using the resting state results as the control, we specifically attempted to answer the following three questions: 1) how do QPP waveforms differ between systematic and random visual stimulation conditions? 2) how do the different visual stimulation sequences impact the frequency and/or intervals of consecutive QPPs? and 3) how do the different QPP phases prior to the stimulus modulate the subsequent visually evoked BOLD responses in different visual stimulation conditions?

## Methods

### 2.1. Data acquisition and preprocessing

Functional MRI brain images of fourteen young adults (8 women, 6 men) in the Atlanta area participated in this experiment (mean age = 19.8±1.7 yro; range [18-24 yro]). The fMRI scanning was performed at the Center for Advanced Brain Imaging (CABI) in Atlanta on a 3T Siemens Trio scanner with a 12-channel radio frequency coil. For each subject, 7 complete gradient-echo echoplanar imaging (EPI) scans were acquired followed by an anatomical T1 image (MPRAGE; TE=3.98ms, flip angle=9°, matrix 256×256 (RO×PE), Slice thickness=1.0 mm,176 slices and voxel size 1×1×1mm^3^). Each EPI scan has 870 timepoints, i.e., *N_x_* = 870 with the sampling rate, *TR* = 0.7s for a duration of 10min and 9s. Other acquisition parameters of EPI scans include TE=30ms, flip angle=90°, Matrix 64×64 (RO×PE), Slice thickness=3.0mm with 22 slices and voxel size 3.4375×3.4375×3mm^3^, Multiband factor=2, Echo spacing = 0.51ms, and BW = 30.637Hz/Px.

Due to the short TR (TR=0.7s) of the single-shot gradient echo-planar imaging (EPI), it is not possible to scan the entire brain. Therefore, certain brain regions are excluded from the EPI scan. As the effects of flickering checkerboard on the visual cortex and visual processing have been extensively studied in the past (Dale & Buckner, 1997; Engel et al., 1997; Schwartz et al., 2005; Tootell et al., 1998), the current study focused on how this visual stimulation, known to activate or deactivate various regions across the whole brain (Jorge et al., 2018), affected the remaining brain regions, such as the default and task-positive networks. Hence, the orbital frontal cortex, temporal pole, dorsal motor areas, and occipital lobe were excluded from each EPI imaging scan, and the corresponding regions were identified using the Schaefer-Yeo Atlas (Schaefer et al., 2018) in the final preprocessing step.

Each of the seven EPI scans for each subject fell into one of three distinct experimental conditions. For all subjects, a resting state scan lasting 10.15 minutes (870TRs) was the first functional scan collected. During the resting scan, subjects were told to stay awake and remain still while staring at a fixation cross. After the completion of the resting state scan, six visual stimulation EPI scans were collected using two visual stimulation conditions. During both conditions subjects were told to focus on a red fixation cross at the center of the projection screen and that on occasion a flashing checkerboard would appear in the background. For both conditions, the flashing periods were comprised of a black and white checkerboard pattern that inverted every 5 refresh frames (60Hz) for a period of 2.1s (3TRs). The red fixation cross remained at the center of the screen during the checkerboard periods. Half of the scans used a *systematic stimulation* sequence. That is, a flashing checkerboard stimulus appeared for 2.1s (3TRs) every 20.3s (29TRs). The other three scans used a *random stimulation* sequence. That is, the flashing checkerboard stimulus appeared randomly at every 13.3s~61.6s (19~88 TRs, average arrival time 19.25s+/−6.34s). For the systematic condition, the stimulus interval of 20.3s ensures at most one QPP occurring during or after each stimulus onset. For the random condition, multiple QPPs could occur between some of the long stimulus intervals. For both visual stimulations, the range of stimulus intervals was selected to warrant a total of 30 stimulation onsets during each EPI scan, which allows us to compare the interaction between each stimulation and ongoing QPP between the two visual conditions. The two types of stimulation sequences alternated in an ABABAB order, with the order counterbalanced between subjects. An illustrative example of these two different visual stimulation sequences is provided in Fig. S1. Between each EPI scan, there was a roughly 30s-time gap, during which we told the subjects to rest their eyes and remain still, inquired on their wakefulness during the preceding scan and informed them of the time remaining until the end of the scanning session.

The acquired fMRI data were preprocessed by an automated pipeline based around SPM12 (https://www.fil.ion.ucl.ac.uk/spm/software/spm12/), FSL (Jenkinson et al., 2012), and AFNI (Cox, 1996; Cox & Hyde, 1997). First, the anatomical T1 image was spatially normalized to the 2 mm Montreal Neurological Institute (MNI) atlas. This step includes an image reorientation to the MNI space using FSL, a bias-field correction using FEAT (Y. Zhang et al., 2001), and the SPM segmentation model, which performed the tissue segmentation of grey matter, white matter, cerebrospinal fluid (CSF), bone, soft tissue and air/background of gray matter, and the spatial normalization of these segmented tissues. The binary mask of the white matter, CSF, and the whole brain (gray matter, white matter, and CSF) were obtained by thresholding at the top 70% of these normalized tissues.

Next, the functional EPI timeseries were preprocessed following procedures as described in (Abbas, Belloy, et al., 2019). Specifically, the following six steps were performed. First, in order to normalize all scans of each subject to the same template, all seven EPI scans of each subject were concatenated. Second, the concatenated EPI data was reoriented (FSL), realigned (SPM12), and normalized to the MNI atlas based on the estimates of the SPM segmentation model from the anatomical data preprocessing. In parallel, the motion parameters for the concatenated EPI data, including the framewise displacement (FD), were also estimated using MCFLIRT (FSL). Here, the FD is estimated by the relative root-mean-square movement (Jenkinson et al., 2002) in order to examine the in-scan head motions in step six. Third, the normalized EPI data were spatially smoothed with a Gaussian kernel of 4 mm (SPM smooth). Fourth, the concatenated EPI data was split back to each scan. Fifth, the EPI data of each scan was temporally filtered at a bandwidth of 0.01Hz~0.1Hz (AFNI 3dBandpass), and was further regressed by the mean signals extracted from the white matter and CSF masks. Sixth, for each scan, the head motions were examined by the FD following the criteria described in (Yousefi et al., 2018). Specifically, scans with low to moderate levels of motion (i.e., mean FD<0.2mm and with a temporal ratio of FD>0.2mm smaller than 40%) were included in our analysis (see Fig. S2 for more details), because the low to moderate levels of motion was found to have minimal impact on the QPP being detected (Yousefi et al., 2018). Note that because the preprocessing procedures described above have demonstrated success in detecting QPPs from resting as well as task-evoked human brains (Abbas, Belloy, et al., 2019), additional preprocessing procedures such as motion parameter regressions and volume scrubbing were not performed.

Finally, the preprocessed EPI timeseries were extracted from the brain parcels provided by the Schaefer-Yeo Atlas (Schaefer et al., 2018) (<u>webpage</u>) and then z-scored. Due to the incomplete brain coverage of the EPI scans (as described in the 2^nd^ paragraph of this section), only parcels with over 85% coverage across all subjects were selected from the Schaefer-Yeo Atlas. The percentage of EPI scanning coverage of each Schaefer-Yeo parcel of each subject is reported in Fig. S3. Notably, because the primary and the majority of the secondary visual cortex weren’t covered for all EPI scans, the visual network was excluded from the analysis. In addition, because the temporal lobe was not covered and most of the remaining parcels in the limbic system have less than 85% coverage, the limbic network was also excluded from the analysis. As a result, EPI timeseries from a total of 193 parcels were extracted, which covered the 5 functional brain networks as described in (Thomas Yeo et al., 2011) including the Somatomotor (SM), Dorsal Attention (DA), Ventral Attention (VA), Frontoparietal (FP) and Default (D) networks.

### 2.2. Quasi-periodic pattern detection and examination

The primary QPP is a phase-locked spatiotemporal pattern detected from the BOLD fluctuations, which repeats over time. This intrinsic dynamic pattern has been found to tie to the infraslow electrical activity (<0.1Hz) (Grooms et al., 2017; Raut et al., 2021; Thompson et al., 2014). In this section, we first describe 1) the detection of the primary QPP and the corresponding QPP with a reverse phase, and then 2) the parameters used for detecting QPPs. Moreover, we describe 3) the rationale and approach of detecting QPPs at a group level, and 4) the procedures for comparing different QPPs in the final paragraph.

#### 2.2.1. Primary QPP, reverse phase QPP, and their occurrences

Primary QPP and its occurrence across the entire timeseries for each experimental condition was detected on the EPI data using the robust QPP detection algorithm described in (Yousefi & Keilholz, 2021). This is a correlation-based and iterative finding algorithm, which identifies similar segments of a functional timecourse and averages them for a representative spatiotemporal template. The algorithm detection process can be summarized by the following six steps. First, an initial segment with a preset window length (WL) was selected at the *i^th^* timepoint (*i* = 1, …, *N_x_ − WL*) of the EPI timeseries of all ROIs. This initial segment has a spatial dimension and a temporal dimension (ROIs × WL) and was used as the reference QPP template for later steps. Second, the reference template was correlated with a segment with the same window length across all ROIs, which was sliding from the 1st to the *N_x_ − WL* timepoints of the timeseries at a step of 1 timepoint, which resulted in a timecourse of sliding correlations. Third, local maxima of this correlation timecourse, which are above a preset positive threshold and also have a minimum distance of WL, were selected as the occurring time of the reference template, and segments with starting points at these local maxima were averaged to obtain an updated template. Fourth, steps 2 and 3 were iterated until the averaged template and the reference converge. Fifth, steps 1-5 were repeated for all *i*s (*i* = 1, …, *N_x_ − WL*), which resulted in totally *N_x_ − WL* sets of results. Notably, the detection process omits the final WL timepoints to avoid QPP finding at the time boundary of different scans. Sixth, the *N_x_ − WL* sets of results were ranked based on the summation of local maxima of the correlation timecourse, and the set of results with the greatest summation was selected as the final solution. This entire process, also summarized in the flowchart (Yousefi & Keilholz, 2021, Fig. S1), produced two major outputs, one is the averaged 2D template of timeseries, which is the primary QPP; the other is the timecourse of sliding correlation, of which selected local maxima are considered as of the occurrence of this QPP. It is worth mentioning that the primary QPP in resting humans displays a sinusoidal waveform (Abbas, Belloy, et al., 2019; Belloy, Shah, et al., 2018; Yousefi et al., 2018; Yousefi & Keilholz, 2021). Specifically, the primary QPP in resting human brains is half-wave symmetric, comprising nearly identical half-cycles with opposite polarities (Yousefi & Keilholz, 2021, Fig. S8b).

Each primary QPP is paired with a corresponding QPP in the reverse phase, known as the reverse phase QPP^1^. While the primary QPP is obtained by averaging the segments that start from the selected local maxima of the correlation timecourse, the reverse phase QPP can be obtained by averaging the segments starting from the selected local minima of the same correlation timecourse. These selected local minima were required to be separated by at least WL and have a negative magnitude below a predetermined threshold. In the resting dataset of the Human Connectome Project (HCP), the primary QPP detected in concatenated scans of a subject may begin from a positive magnitude (like the sine wave) or a negative magnitude (like the −sine wave). Remarkably, the primary QPP with a “-sine” waveform is highly similar (Pearson correlation r>0.88, p-value<0.01) to the reversed phase QPP associated with a primary QPP with a “sine” waveform (Yousefi & Keilholz, 2021, Fig. S25-S26, Video 3). Hence, both the primary QPP and its reverse phase QPP have been utilized in the primary QPP analysis for studying resting-state populations. However, when analyzing the occurrences of QPPs for the resting populations (see Section 2.3), we only considered the occurrences of the primary QPPs. This is because the QPP correlation timecourse was selected based on the summation of the local maxima, which were limited to the primary QPP only, and not on its reverse phase counterpart or the local minima.

#### 2.2.2. Parameters for QPP detection

The QPP window length was selected based on the duration of QPP templates observed in previous studies (Majeed et al., 2011b; Thompson et al., 2014; Yousefi et al., 2018), and a common window length was chosen for easy comparison across different experimental conditions (e.g., for the correlation calculation described in the final paragraph). Because the duration of QPP lasts for approximately 20s in both resting (Majeed et al., 2011b; Thompson et al., 2014; Yousefi et al., 2018) and evoked human brains (Abbas, Belloy, et al., 2019), QPP window lengths (WLs) ranging from 17.5s to 24.5s (WL=25TRs~35TRs) were explored. The final window length was determined by identifying the point at which increasing the window length would no longer change the appearance of the QPP across all experimental conditions. This ensured that the selected window length was appropriate for detecting the primary QPP in all experimental conditions. In addition, the positive correlation thresholds and maximum iterations were selected based on previous studies (Abbas, Bassil, et al., 2019; Abbas, Belloy, et al., 2019; Belloy, Naeyaert, et al., 2018; Belloy, Shah, et al., 2018; Majeed et al., 2011a; Yousefi et al., 2018). In particular, a positive QPP correlation threshold of 0.1 for the first three iterations and 0.2 for subsequent iterations was selected with a maximum of 20 iterations. A negative correlation threshold of −0.2 was set for detecting the reverse phase QPP. The above parameters have been shown to be effective in detecting primary QPPs across various experimental conditions in humans (e.g., TR=0.3s~2s, resting state, N-back task, and disease models) (Abbas, Bassil, et al., 2019; Abbas, Majeed et al., 2011a; Yousefi et al., 2018).

#### 2.2.3. Group QPP analysis for each experimental condition

Group QPP analysis is a common approach for comparing primary QPPs across different experimental conditions or populations (Abbas, Bassil, et al., 2019; Abbas, Belloy, et al., 2019; Majeed et al., 2011a). In this approach, all EPI scans from each experimental condition or population are concatenated into a single timeseries, which is then subjected to detection of the primary QPP. By comparing the primary QPPs across different groups, researchers can gain insights into the differences or similarities in brain dynamics between different conditions (Abbas, Bassil, et al., 2019; Abbas, Belloy, et al., 2019; Majeed et al., 2011a). Recently, a subject-level QPP analysis was employed on the high-quality resting-state HCP dataset, in which 4 EPI scans of each subject were concatenated (resulting in a 57.6-min concatenated EPI data) for the QPP detection (Yousefi & Keilholz, 2021). This approach could reveal inter-subject variabilities within one experimental condition.

In our study, because we aim to investigate how different sequences of visual stimulation (*systematic* or *random*) influence spatiotemporal brain dynamics, and because we have far less EPI data for each experimental condition for a subject-level analysis (i.e., a total of 10.15-min EPI recording for resting and 30.45-min EPI recording for the systematic and for the random conditions), we have designed a group QPP analysis with repeating groups to assess the variabilities within each visual condition. Specifically, the extracted EPI timeseries from each scan were concatenated across all 14 subjects, which results in 1 resting group, 3 repeating groups with systematic stimuli and 3 repeating groups with random stimuli. Each group includes a 142.1-min concatenated EPI data, which according to our previous study (Yousefi & Keilholz, 2021) is sufficient for providing reproducible results under one experimental condition. The three repeating groups with visual stimulations will be used to test the variabilities and reproducibility within each visual condition. In the remainder of the paper, we refer to this group EPI timeseries as the resting, the systematic 1, 2, or 3, and the random 1, 2, or 3.

For the resting state condition, the group primary QPP (*QPPrest*), and its reverse phase QPP (*QPPrest*-), as well as their occurrences were obtained from the detection algorithm. For the systematic condition, timeseries of all 3 repeating groups were first concatenated for generating a *group average QPP* (*QPPsys*) and the correlation timecourse across all 3 groups. Then, the QPP for each group (systematic 1, 2, and 3) was obtained by only averaging the EPI segments starting at the correlation local maxima in each group (*QPPsys_i_* for *i* = 1, .., 3). Similarly, the group average QPP (*QPPrand*) and three group QPPs (*QPPrand_i_* for *i* = 1,.., 3) were obtained for the random stimulation data. The systematic and random *group average QPPs* have the following relationships: 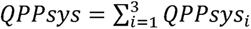 and 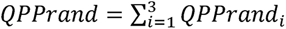. In addition, the reverse phase QPP was also obtained for each visual condition, denoted by *QPPsys*─ and *QPPrand* ─. We refer to this set of results as *group analytical results*. An illustration of these results and the process that arrived at them are shown in Fig. S5. To further test the reproducibility of the results, we also performed an *independent group analysis*, which detected the QPP independently for each task group, which obtained 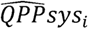 and 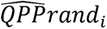 for *i* = 1,.., 3. The data used for the *group average analysis* and the *independent group analysis* was shown in Table S1.

#### 2.2.4. Comparison between two primary QPPs

Comparing multiple primary group QPPs is a common approach to examining spatiotemporal brain dynamics across different experimental conditions or populations (Abbas, Bassil, et al., 2019; Abbas, Belloy, et al., 2019; Majeed et al., 2011a). Here, we aimed to assess the differences between the primary QPP waveforms in the systematic and random visual conditions, however, we found that *QPPsys* and *QPPrand* have opposite phases. It’s worth noting that the prior study (Yousefi & Keilholz, 2021) discovered that both *QPPrest* and *QPPrest* - were suitable for primary QPP analysis in the resting-state. To facilitate comparison, we measured the differentiation between the QPPs of each visual condition and the resting QPP with the same phase. The QPP observed during the visual stimulation was considered the *empirical result*, while the resting state QPP used for comparison was referred to as the *null model*.

More specifically, we performed the assessment in three steps. Firstly, we calculated the correlation coefficients between the empirical (*QPPsys* or *QPPrand*) and the null model (*QPPrest* or *QPPrest*-). Secondly, we conducted a z-test to determine if the correlation of the two visual conditions differed significantly from each other. Specifically, the Fisher z-transformed correlation coefficient was computed from each condition and then the z-score of their differences was computed with sample normalization (Diedenhofen & Musch, 2015). The significance of the difference was also tested at the average level of the three groups. Finally, we examined the distinctions in the entrained QPP waveforms from the following four aspects: 1) the phase shift from the null, 2) the amplitude changes in percentage from the null, 3) the vertical shift in the percentage of the amplitude of the null, and 4) the percentage changes in “peak-life” of the positive and negative peaks from the rest (see Fig. 1A). Because the QPP of parcels in the same brain network was found to share the same waveform (Yousefi et al., 2018; Yousefi & Keilholz, 2021), the waveform examination was conducted at the network level, i.e., the network QPP obtained by averaging the QPP of parcels in each of the five networks described in the second to the last paragraph of Section 2.1. More specifically, the phase shift was estimated using (Zhivomirov, 2022) which implements the algorithm in (Sedlacek & Krumpholc, 2005). The “peak-life”, in particular, is defined as the entire period that the waveform stays from its peak to half of its peak. A 2-way ANOVA and multiple comparison tests were used to compar the two visual conditions for each waveform characteristic across all networks.

**Figure 1.**
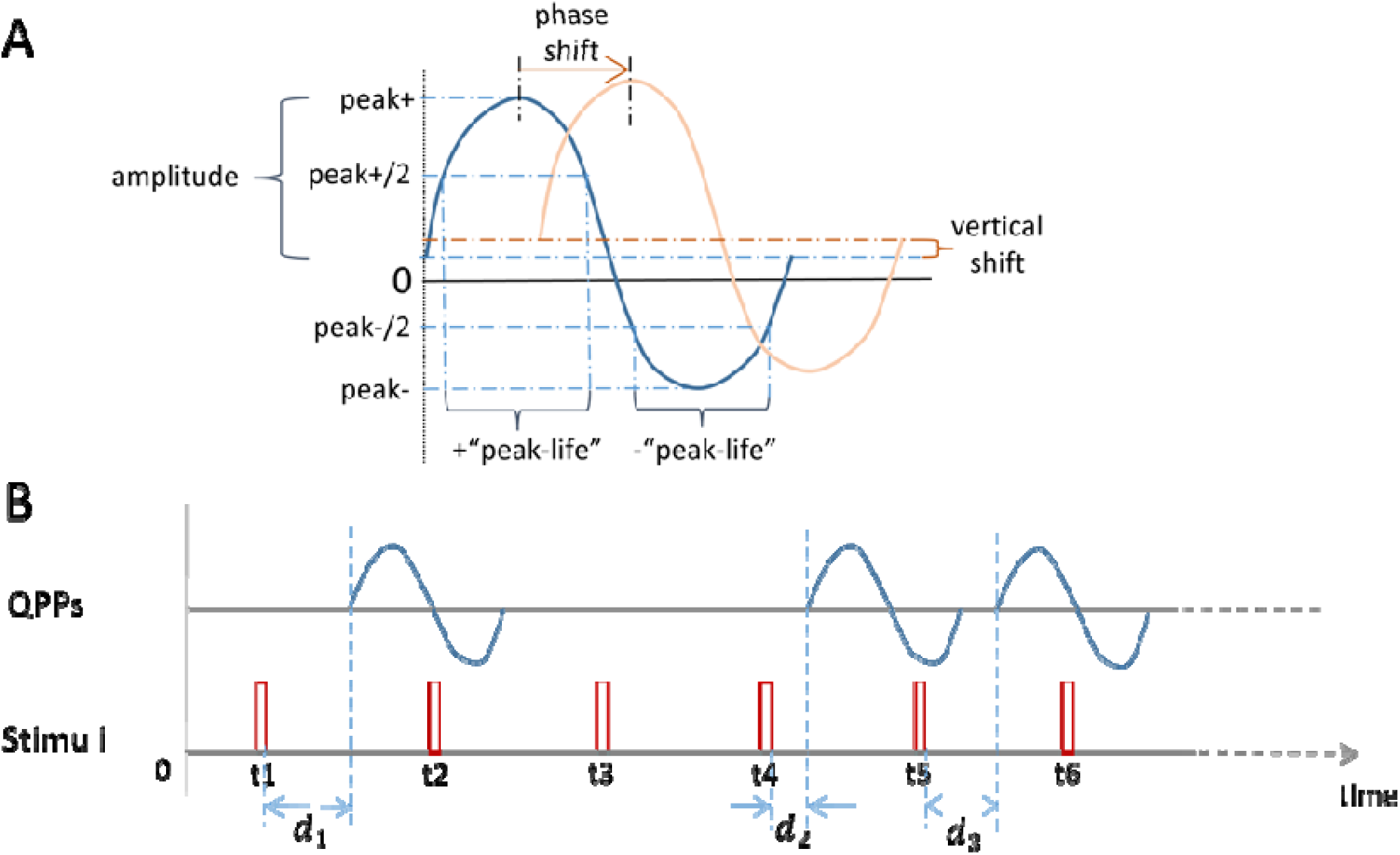
QPP waveform and time delays of QPPs occurrence followed by stimuli. (A) characteristics that describe QPP waveform distinctions, which include the phase and vertical shifts, the amplitude changes, and distinctions in the “peak-life” of the positive and negative peaks. (B) time delay of each ongoing QPP post the stimulus onset, denoted by *d_i_*. Only nonzero *d_i_*’s were included in the analysis.

### 2.3. Entrained QPP occurrence analysis

The study also investigated the association between the onset of visual stimuli and the occurrence of QPPs. This analysis utilized the primary QPP occurrences obtained from both the group average and independent group results (see Table S1). The investigation aimed to answer two questions. The first question examined whether the type of visual stimulation sequence, i.e., systematic or random influences the frequency and intervals of consecutive QPPs. The second question aimed to investigate whether the onset of stimulation in either visual condition affects the timing of ongoing QPPs (i.e., advance or delay the onset of successive QPPs).

To address the first question, we calculated the frequency of QPP occurrence over time for each group. This was done by dividing the total number of QPP occurrences by the number of timepoints in the 14 concatenated scans. The resulting frequency was then averaged across the repeating groups and compared between the three experimental conditions. Additionally, the intervals of consecutive QPPs were also computed, and the mean of the intervals was compared between the systematic and random visual conditions. A t-test was then performed to determine if the measured mean difference between the two conditions was significantly greater or smaller than zero.

To investigate the second question, we calculated the time delay of QPP occurrence after each stimulus (Fig. 1B) and formulated two hypothesis tests, one for each visual condition. The null hypothesis was that the visual stimuli did not affect (i.e., delay or advance) the timing of ongoing QPPs, while the alternative hypothesis was that they did. For each visual condition, the null model assumed that the timing of QPP was the same as that observed during the resting state. The null distribution of QPP delay was constructed by comparing the correlation timecourse of resting QPP with the stimulation sequence and calculating the time delay of QPP occurrence following each visual stimulus. Different null distributions were generated for each visual condition as the two types of stimulation sequences had different stimulus onset times, which introduced different cutoff values for QPP delays (i.e., <29TRs for the systematic condition and <88TRs for the random condition). An example of null and empirical model of QPP delay is shown in Fig. S9. Finally, for each hypothesis test, a two-sample t-test was used to determine if the mean of the empirical result significantly differed from the null model.

### 2.4. QPP phase-dependent BOLD stimulus response analysis

We investigated how QPPs affect the BOLD response to different stimulation sequences from the following two aspects. First, we examined the distinction in the BOLD-stimulated responses between two visual conditions. We focused on the peak value of each stimulus response, which was measured by the shaded area in Fig. 2A. More specifically, we measured the BOLD magnitude in the time window of 4.2s~8.4s (equivalent to 6TR~12TR, a total of 7 timepoints) after stimulus onset. Then, we subtracted the BOLD magnitude at the last timepoint pre-stimulus from this measurement. Finally, we summed this difference over all 7 timepoints to obtain the peak value (Fig. 2A). This measurement is similar to the calculation described in (He, 2013; Huang et al., 2017), and is due to the fact that the hemodynamic response peaks about 6s post the stimulation onset and has a bandwidth of 4s (Wald & Polimeni, 2015). Means of systematic and random stimulus response as well as the systematic-random contrast were computed across the entire stimulation sequence. Parcels with significant systematic-random contrast were identified by p-value<0.05 (equivalently |zscore|>1.96) after z-scoring the averaged contrasts among all 193 parcels.

**Figure 2.**
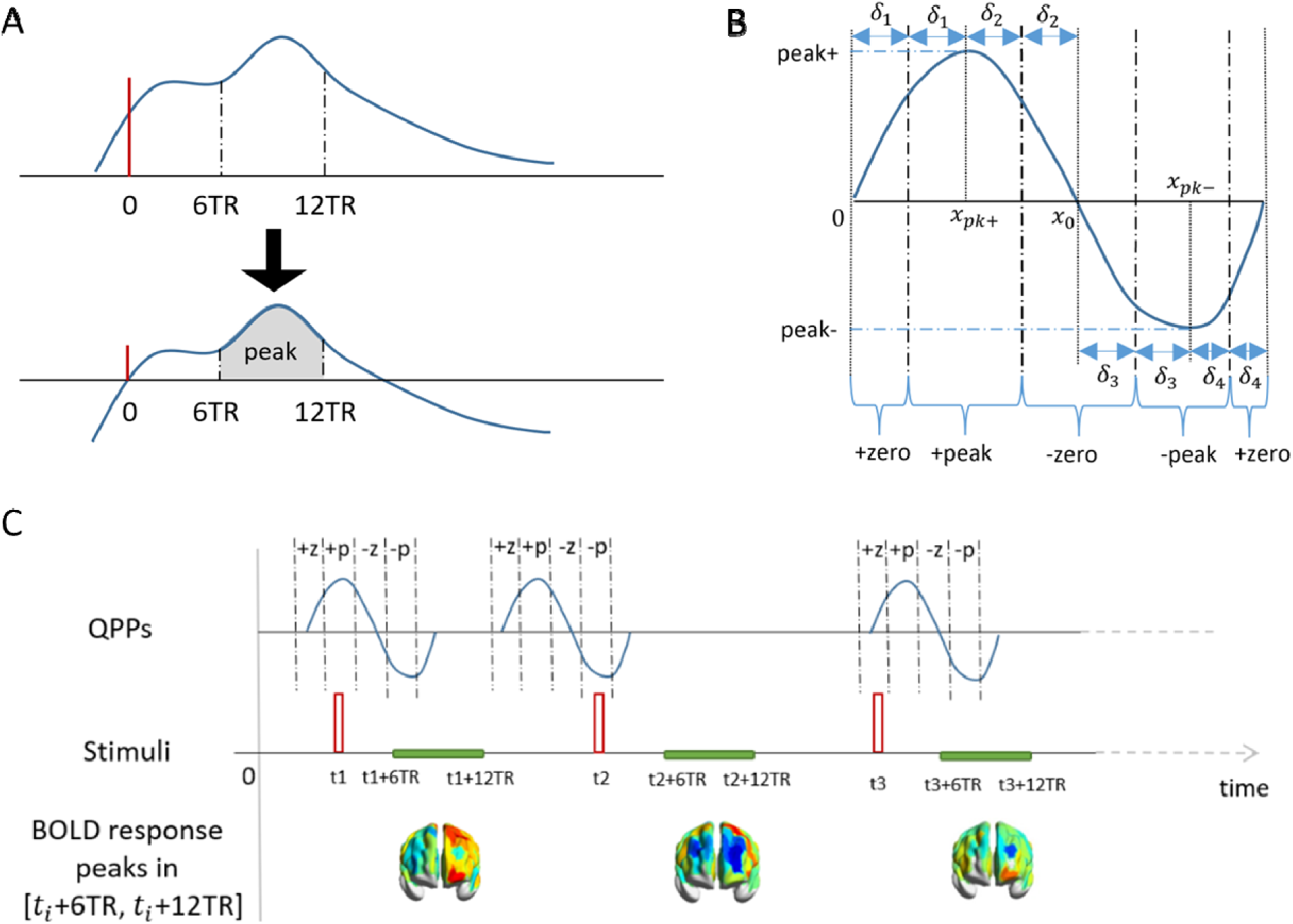
Dependence of the BOLD response to visual stimulation and QPP phase. (A) determination of peak value (shaded area) of evoked BOLD responses. The red vertical line depicts the onset of visual stimulation. The BOLD magnitudes in [6TR, 12TR] were subtracted by the BOLD magnitude at the stimulus onset. The gray area under the bottom curve depicts the peak value of the stimulus response. (B) illustration of four-phase zones of a QPP. Let, and be th timepoint of the positive peak, negative peak, and the midpoint where the QPP wave across the zero, respectively. Based on these timepoints, four pairs of intervals can be determined as –––, ––––, ––––, and ––––, where WL is the QPP window length. Then, the four phases +zero, +peak, −zero, −peak, have the following zones [0, [ ], [ ,], [ , ], [ , ]. (C) BOLD stimulus with stimulus onset overlaps ongoing QPPs at different phases.

To investigate the effect of visually evoked BOLD responses on overlapping QPP at different phases, we divided the QPP of each parcel into four distinct phase zones, referred to as “+zero”, “+peak”, “-zero”, and “-peak”. This division was achieved through a 4-step process. Firstly, the timepoints of the positive and negative peaks of the QPP wave, as well as the midpoints where the wave crossed zero, were determined for each parcel. Secondly, we identified four pairs of non-overlapping time intervals (δ*_i_* for *i* = 1,2,3,4) covering the entire window length [0, *PL*]. Thirdly, we defined the “+peak” (“-peak”) phase zone as the interval(s) stepping away from the positive (negative) peak within the corresponding *o*_i_. Finally, the “+zero” (“-zero”) zone was identified as the non-overlapping interval(s) containing the QPP across zero with an uprising (a down-falling) trend. The four-phase zones for an exemplary QPP waveform are illustrated in Fig. 2B and the BOLD stimulus responses overlap different phases of the ongoing QPP are illustrated in Fig. 2C.

The BOLD responses to the stimuli that coincide with each phase of QPP (*QPPsys*, *QPPrand*) and of its reverse-phase counterpart (*QPPsys*-, *QPPrand*-) were averaged across the stimuli. Additionally, a control group was included by averaging the BOLD responses to stimuli that did not overlap an ongoing QPP. The resulting averaged BOLD responses for each phase were then compared across brain regions (parcels) and between the two types of stimulation sequences (systematic and random), as well as between the empirical and null results.

## Results

### 3.1. Differences in group-level QPPs across systemic and random stimulations

First of all, the group average QPP affected by the systematic stimulations (in comparison to the random stimulations) appeared to be more distinct from the resting state. Differences in group average QPPs for each visual condition in comparison to the resting QPP were shown both among all parcels (Fig. 3A) and for each network (Fig. 3B). Numerically, the group average resting QPP has a significantly lower correlation to *QPPsys* than to *QPPrand* (i.e., z-score=-36.76, p-value<0.01), as the fisher z-transformed correlation between *QPPrest* and *QPPsys* is 1.003 (Pearson-correlation r=0.763, p-value<0.01), and it is 1.673 (Pearson-correlation r=0.932, p-value<0.01) between *QPPrest*- and *QPPrand*. This significantly lower correlation value also appears in the between-group calculations (Table S2, z-score=-32.402, p-value<0.05), as well as in the average of independent groups analysis (Fig. S7, z-score=-12.396, p-value<0.05). The group average *QPPsys*- and *QPPrand*- are shown in Fig. S4. The resting QPPs demonstrate (anti-)correlations between different networks that are similar to the previous findings on the HCP dataset (see Fig. S6). The positive and negative phases of both *QPPrest* and *QPPrest*-appeared to have same duration, which is also consistent with the previous findings (Yousefi & Keilholz, 2021). QPPs detected for each independent group are shown in Fig. S7. For each experimental condition, the QPP window length of 21.7s (WL=31TRs) was selected following the procedure as described in Section 2.2.

**Figure 3.**
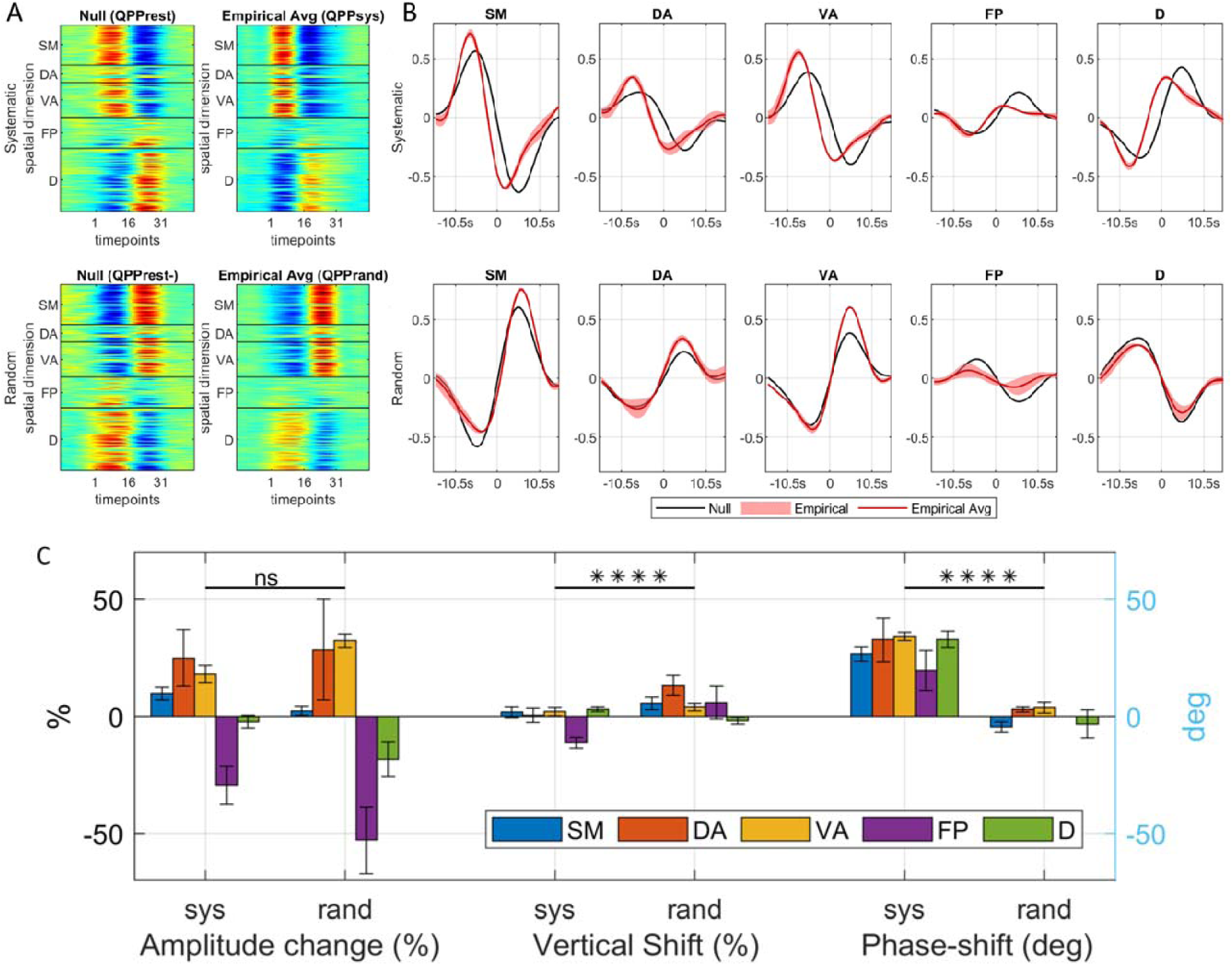
Group average QPP for resting, systematic, and random visual stimuli. (A) the global spatiotemporal QPP across different experimental conditions. For each visual condition, both empirical and null QPPs are shown. These QPPs were simultaneously detected from all 193 parcels, covering five networks, somatomotor (SM), dorsal attention (DA), ventral attention (VA), frontoparietal (FP), and default (D). The y-axis of each pattern corresponds to the spatial dimension, while the x-axis corresponds to the temporal dimension. (B) network QPP averaged among parcels in each of the five networks. Both the null and empirical QPP curves are displayed in each plot, with alignment at the first timepoint. To emphasize the changes in waveform of the empirical QPP for each network, the timepoint where the magnitude of the null QPP curve crosses zero was selected as th reference point (0s). (C) Bar plots of the average changes in three characteristics of the QPP waveform, including the amplitude changes (%), vertical shift (%), and phase shift (deg), as perturbed by the visual stimulation (empirical) when compared to the resting QPP (null). In the case of the frontoparietal network, because the amplitude of is attenuated by over 50% and is almost close to zeros, the phase shift cannot be appropriately determined, and hence a ‘NA’ is reported. The significance level of the multiple comparison test between the two visual conditions is denoted above each characteristic, with ‘ns’ and ‘****’ representing Bonferroni corrected p-values greater than 0.05 and less than 1e-4, respectively. Bar plots of the average changes in all five waveform characteristics, including the + and −“peak-life” changes (%) and the above three characteristics are shown together with the numerical values for these changes and the multiple comparison test results in Table S3.

Secondly, systematic stimulations versus random stimulations appear to affect the network QPP waveforms in different ways. Compared to the random stimuli, the systematic stimuli have a more significant effect (Bonferroni corrected p-value<1e-4) on the phase of QPPs. On the other hand, the random stimuli have a more significant effect (Bonferroni corrected p-value<1e-4) on the magnitude of QPPs (Fig 3C and Table S3). Specifically, in the random condition, the magnitude of QPP shifts more positively in the somatomotor and the three task-positive networks, but shifts more negatively in the default network compared to the systematic condition. This QPP vertical shift is also combined with changes in amplitude. For example, QPP amplitudes increase more in dorsal and ventral attention and decrease more in frontoparietal and default for the random than the systematic condition. In addition, the phase shift affected by the systematic stimuli (an average of 29.18 degrees) is greater than the one affected by the random stimuli (an average of |3.65| degrees). This greater phase shift by systematic stimulations is also reflected as a squeezed +peak in the task-positive networks for both stimulation sequences. For example, the QPP wave’s +“peak-life” in dorsal attention, ventral attention, and frontoparietal networks is shortened by 35.41% on average by the systematic stimuli but only shortened by 18.48% on average by the random stimuli.

### 3.2. Effect of visual stimulation on the incidence of QPPs

On average, the systematic visual condition had a higher frequency of QPP occurrences compared to the random condition, while the resting state was in between (see Table 2 for the group average frequency). This finding is also supported by the shorter QPP intervals observed in the systematic condition compared to the random condition, as illustrated in Fig. 4A. In particular, as shown in Fig. 4A, the mean of the random condition QPP intervals is significantly greater than the systematic condition (i.e., p-value=0.002<1%). In the independent group analysis, the mean of QPP intervals for the three random groups is also greater than for the three systematic groups (see Fig. S8, i.e., p-value=0.007<5%).

**Figure 4.**
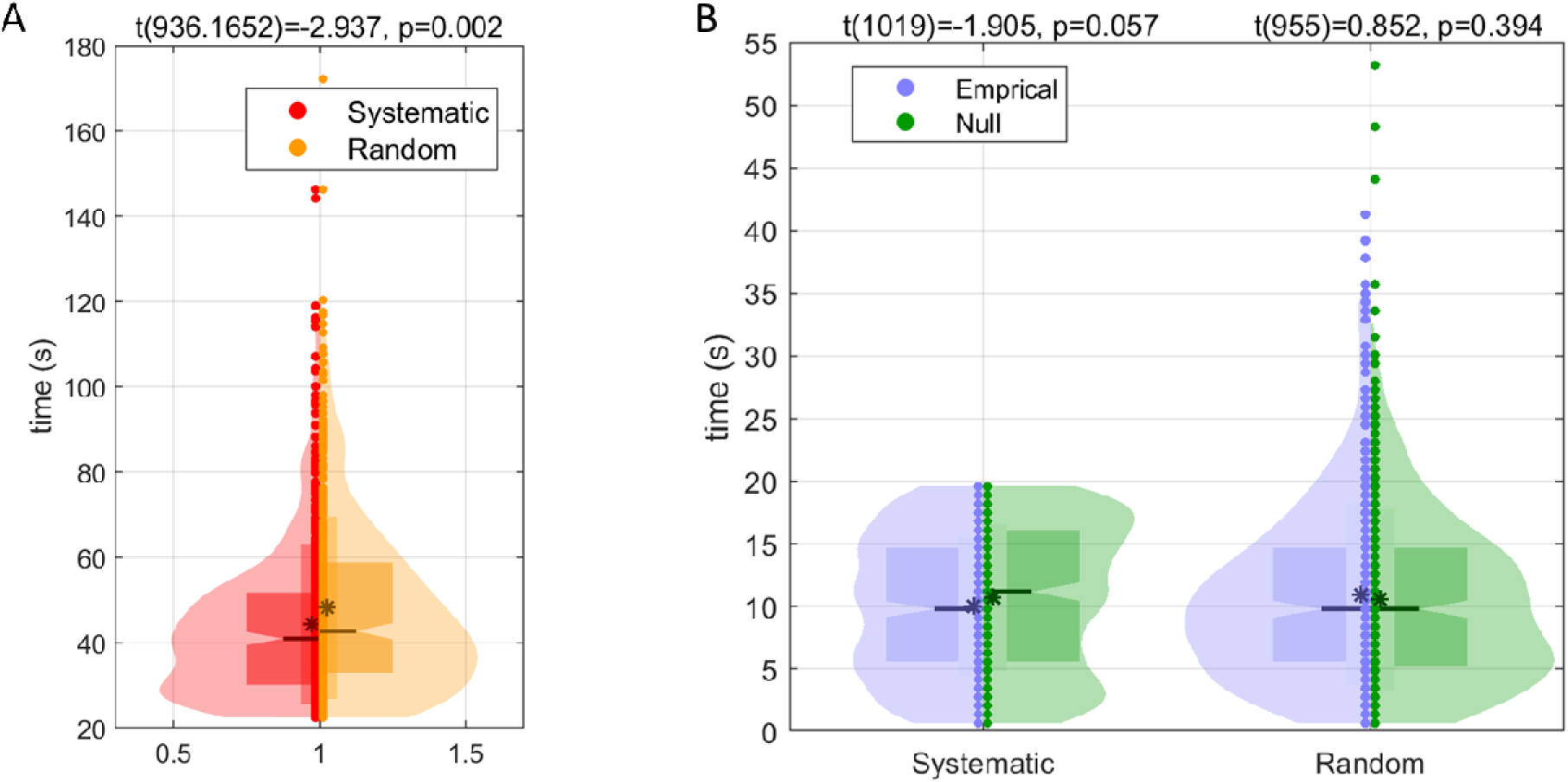
Violin plot of QPP intervals and QPP time delays. (A) Distribution of QPP intervals for all groups for each visual condition. Here, the QPP intervals are contrasted between the two stimulation sequences. (B) Distribution of QPP time delay followed by visual stimuli for all groups in systematic stimulation sequences. Here, the distribution of QPP time delay in each visual condition is contrasted to its null model (derived from the resting data as described in Section 2.3). The t-test statistics, degree of freedom (noted as t(df)), and p-values are reported above each pair of violin distributions. Please refer to the right panel of Figure S8 for a detailed explanation of the violin plot.

**Table 2.** Frequency of QPP occurrences for each group. For each experimental condition, the group average result is averaged by the frequency of the group average QPP that occurred in each group, whereas the independent group result is averaged by the frequency of QPP that is independently detected in each group. The cell with a greater average has a darker color.

|  | Group average results | Independent group results |
| --- | --- | --- |
| Resting | 1.46% | 1.46% |
| Systematic | 1.47+/-0.03% | 1.55+/-0.03% |
| Random | 1.41+/-0.07% | 1.45+/-0.05% |

On the other hand, the QPP time delay in each visual condition exhibits no significant difference from its null model (see Fig. 4B). In Fig. 4B, the probability density function of the time interval between the stimulus and the onset of a subsequent QPP was shown. These QPP delays were compared to the null model (computed from the resting data as described in Section 2.3) for each visual stimulation condition. Numerically, despite the slight difference in the mean of QPP delays for each condition, neither difference is significant (i.e., the systematic condition has p-value=0.057>5%, and the random condition has p-value=0.394>5%, and also see Fig. S10 for the independent group results).

### 3.3. QPP phase dependence of BOLD response to visual stimulation

The systematic versus the random visual stimulations appear to evoke different BOLD responses in several brain regions. In particular, results (Fig. 5 and Fig. S11) suggest that both types of visual stimulation sequences activated the prefrontal lobe. However, the temporoparietal junctions were activated by the systematic stimuli but were deactivated by the random stimuli, whereas the middle frontal gyrus was deactivated by systematic stimuli but was activated by random stimuli. Additionally, systematic stimuli significantly deactivated the ventral regions in the posterior cingulate cortex (PCC)-precentral gyrus of the posteromedial cortex whereas the random stimuli significantly deactivated the default regions in PCC-precuneus.

**Figure 5.**
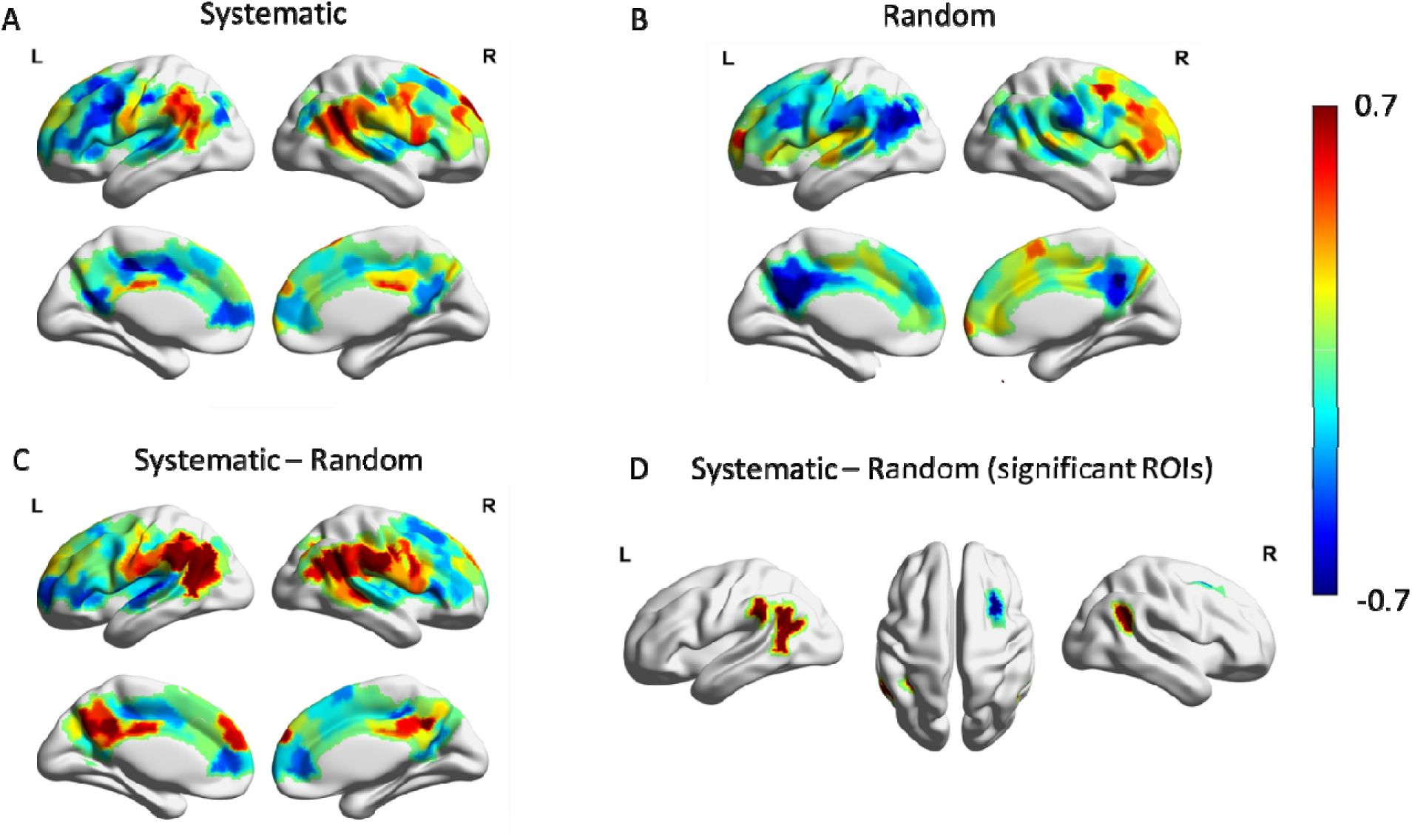
Averaged BOLD peak values in response to systematic and random visual stimulation. See Fig. 2A for the calculation of a stimulated BOLD peak value. (A) The average of BOLD peak values in response to the systematic stimuli. (B) The average of BOLD peaks in response to random stimuli. Parcels with significant (|z-score|>1.96) BOLD peak response for systematic and for random are shown in Fig. S11. (C) The average of systematic-random peak contrast for all selected ROIs. (D) The average of systematic-random peak contrast parcels with significant averaged contrast (|z-score|>1.96). Note that the light grey areas in the brain maps are non-covered regions (see Fig. S3 for the covered and non-covered brain regions in the analysis).

More specifically, six brain parcels demonstrated significant averaged systematic-random contrast (p-value<0.05, as shown in Fig. 5, in which the averaged peak value of BOLD responses for the two visual conditions, as well as for their contrast are demonstrated). Five of them have positive contrast values and all lie in the bilateral temporoparietal junctions spanning across the ventral attention and default networks. On average, these parcels have strong positive BOLD responses to systematic stimuli (e.g., among the top 40% parcels with positive mean of BOLD peaks), but have strong negative responses to random stimuli (e.g., among the bottom 30.8% parcels with negative mean of BOLD peaks). On the other hand, one brain parcel with significant negative systematic-random contrast (zscore<-1.96) is located at the right dorsolateral prefrontal cortex (in the frontoparietal network). On average, this parcel has strong negative BOLD responses to systematic stimuli (i.e., at the bottom 19.44% of parcels with negative mean peak values), but has strong positive BOLD responses to random stimuli (i.e., at the top 6.58% of all positive mean peaks). Among these six parcels, the three task-negative parcels (the three default network parcels) also demonstrate a much more depressed amplitude in averaged QPPs by random stimulations than by systematic stimulations (Fig. S12), which is consistent with the sign of systematic-random contrast in the BOLD response peaks. However, opposite to the systematic-random contrast in the BOLD response peaks, the three task-positive parcels (the two ventral attention parcels as well as the right dorsolateral prefrontal parcel) demonstrate a much more elevated QPP amplitude by random stimulations (Fig. S12).

In addition, the BOLD responses in task-positive and task-negative networks (including the dorsal attention, ventral attention, frontoparietal, and default networks) as well as in the somatomotor network are found to be dominated by the waveform of the overlapping ongoing QPPs. The averaged BOLD responses to stimuli, which were presented at different phases of QPP for each parcel, are organized in the five networks. There are 70.56% of systematic stimuli and 69.84% of random stimuli overlapping with ongoing QPPs. As shown in Fig. 6 and Fig. S15, for both visual stimulation conditions, the BOLD response to the checkerboard is swamped by the ongoing QPP signals no matter which of the four QPP phases overlap. For example, the stimulation onset in the “+peak” (“-peak”) range will follow by a down tread (uprising) BOLD response. In contrast, when the stimulation onset doesn’t meet the ongoing QPPs, an average with more moderate BOLD responses appeared, which covers 29.44% or 30.16% of the entire systematic or random stimuli. The averaged BOLD responses of the six brain regions with a significant contrast between the systematic and random conditions were also linked to the four QPP phases. As illustrated in Fig. S13, both positive and negative contrast values between systematic and random conditions were primarily related to the comparison of ongoing QPPs in each visual condition.

**Figure 6.**
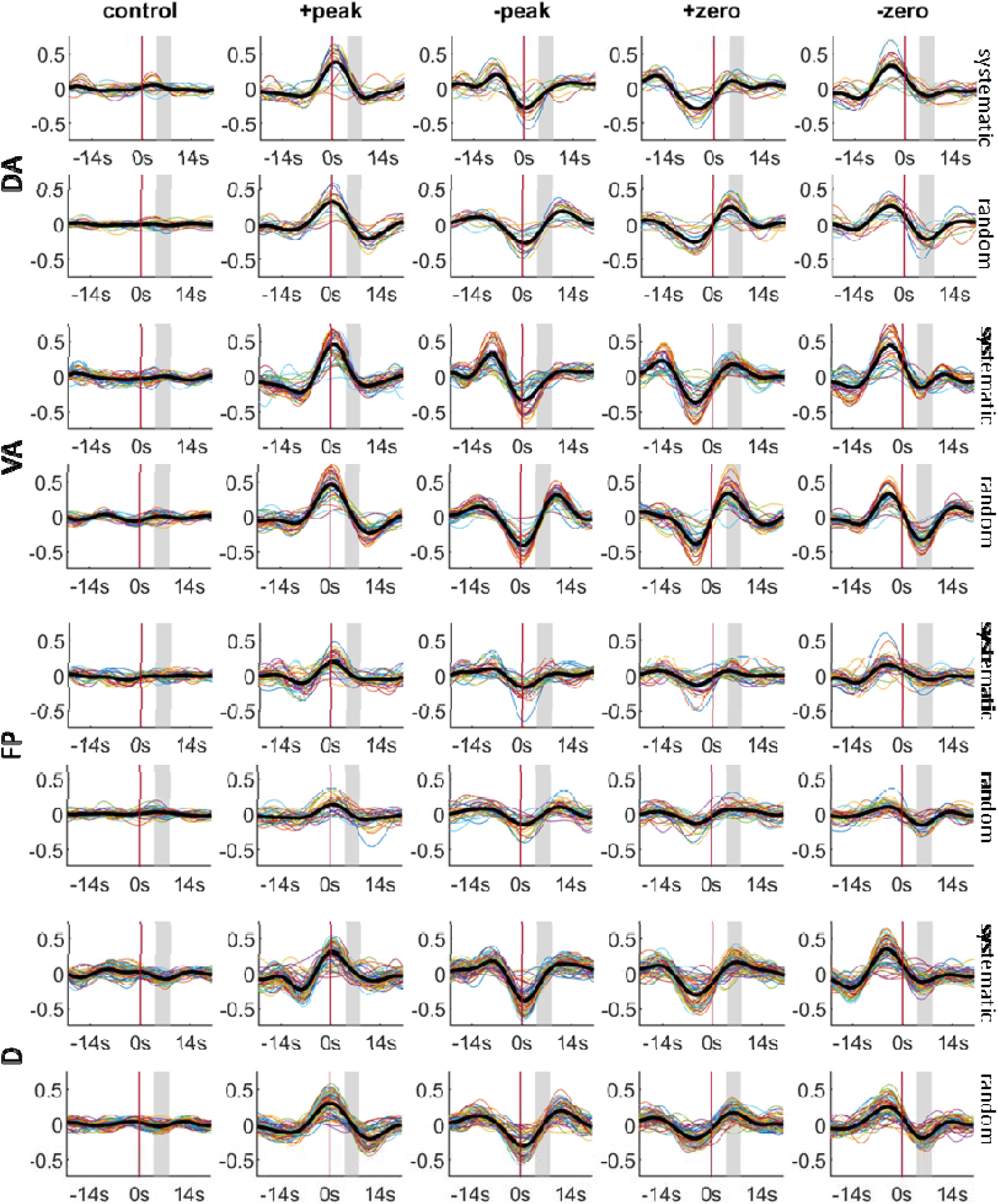
BOLD responses of systematic (upper) and random (bottom) stimulation patterns that are associated with four QPP phases for each of the task-positive and task-negative networks. The control presented in the 1^st^ column includes BOLD responses with no intrinsic primary QPP and the reverse-phase QPP. In each plot, the colorful lines represent the BOLD signal of each parcel within the network, while the bold black line represents the average of all the parcels within the network. The vertical axis represents the magnitude of BOLD response whereas the horizontal axis represents the time interval before and after the stimulus onset at 0s─depicted by the red vertical line. The grey shaded area in each plot depicts the peak range [6TR, 12TR] of the hemodynamic response.

## Discussion

The dynamics of intrinsic brain activity can be captured by several quasi-periodic spatiotemporal patterns (QPPs) (Bolt et al., 2022; Yousefi & Keilholz, 2021). The primary QPP captures the major dynamics of infraslow intrinsic neural activity (Grooms et al., 2017; Thompson et al., 2014), which is known to be involved in attention (Helps et al., 2010; Monto et al., 2008) and arousal (Raut et al., 2021; Sihn & Kim, 2022). The interaction between the ongoing primary QPPs in the brain and visual stimulations was investigated in this study. More specifically, we investigated how different sequences of visual stimuli affect the primary QPP in awake humans, and how spontaneous QPP prior to each stimulus modifies the subsequent visually evoked BOLD response. Two different types of stimulation sequences induced by flickering checkboard were presented to the subjects, a systematic stimulation sequence in which the visual stimulus appeared every 20.3 secs and a random stimulation sequence which has the visual stimulus occurring randomly every 14~62.3 secs. Finally, the results of the two types of stimulation sequences were contrasted to the resting state results, which were then compared with each other.

Due to the limited brain coverage resulting from the use of a short TR and single-shot gradient EPI, this study mainly examined the interaction between visual stimulation and the default, task-positive, and somatomotor networks. This is because flickering checkerboard visual stimuli have been shown to activate and deactivate numerous regions throughout the brain, not just in the visual system (Jorge et al., 2018). In our study, the flickering checkerboard stimuli were designed to have minimal impact on arousal fluctuations, but there may be a slight increase in arousal due to uncertainty (Critchley et al., 2001; Ramsøy et al., 2012; Urai et al., 2017; Zhao et al., 2019) surrounding the random stimulus intervals. While the sequence of visual stimulation is consistent across all systematic scans, it differs across all random scans. Although the designed stimulation sequences may introduce increased variability in arousal levels in the random condition, the standard deviation of QPP correlations in both the group average analysis (see Table S2 bottom) and the independent group analysis (noted in the caption of Fig. S7) indicates that the variability of QPP within the random condition is actually smaller than that within the systematic condition. Thus, we believe that the effect of arousal caused by the uncertain stimulus intervals in random sequences is minimal, or not captured by the QPP. Three central observations of this study are discussed below.

### Sequences of visual stimulation modify the group averaged QPPs

The QPPs during the systematic visual condition are significantly different from the ones during the random visual condition, which is more similar to the resting QPPs. These differences are primarily reflected by a phase modulation. This is consistent with existing literature on higher-frequency activity. For example, in a theta frequency band, the phase of spontaneous oscillations was found to be significantly modulated by only predictable (or attended) but not unpredictable (or unattended) visual stimuli (Busch & VanRullen, 2010). Similarly, in a frequency band of ~8Hz, the phase coherence was found to be strengthened by a systematic visual attentional task (Zareian et al., 2020).

For the random condition, visual stimulation was found to affect the QPP magnitude at a network level. Specifically, the QPP magnitude was much elevated in the ventral and dorsal attention networks but attenuated in the frontoparietal network. The default network QPP was also more attenuated by random stimuli than systematic stimuli. Similar magnitude changes in infraslow dynamics have also been observed in patients with ADHD (Abbas, Bassil, et al., 2019; Helps et al., 2010), suggesting that sustained attention was distracted by random visual stimuli. More specifically, the systematic sequence may entrain intrinsic neural oscillations related to generating expectancies for future events and allocating attention, while a random presentation of stimuli typically involves a longer reaction time and may indicate less sustained attention (e.g., Jones et al., 2002; Lakatos et al., 2008). However, as the study did not include a performance-based measure of attention, these speculations are based on reverse inference and should only serve as a starting point for further research.

### Visual stimulation affects both the frequency of QPP occurrence and the BOLD response to the stimulus

Even though the visual conditions have exactly the same number of stimuli, the systematic stimulation produces more frequent QPPs with significantly shorter consecutive QPP intervals than the random stimulation (Fig 4a). However, neither stimulation sequence significantly perturbs the onset of QPPs. This seems to contradict the previous findings in mice that the primary QPPs are more likely to be triggered at the onset of stimulus (Belloy et al., 2021). There are several possible reasons for this discrepancy. First, a very different visual stimulation sequence was employed in (Belloy et al., 2021). Particularly, a stimulation of “ON” (30s) and “OFF” (60s) cycle that repeats over time includes a flickering light constantly flashing at 4 Hz before becoming silent. Second, anesthetized instead of awake mice were studied in (Belloy et al., 2021), and anesthesia is known to affect infraslow brain dynamics (Pan et al., 2013). Finally, this difference may suggest that the intrinsic QPPs in humans are more robust and less likely to be disrupted by environmental perturbations compared to anesthetized mice.

In addition, we also found that the two visual stimulation sequences evoked distinct patterns in the BOLD response. For example, the bilateral temporoparietal junction (spanning the ventral attention and the default networks) was significantly activated by the systematic condition but not the random condition (Fig. 5D). This is consistent with reports that this region is involved in temporal order judgment (Davis et al., 2009) and lack of predictability in the random condition (Wu et al., 2015). On the other hand, the right dorsolateral prefrontal region (around Brodmann area 9) was strongly activated by random stimulation but not systematic stimulation (Fig. 5D), which contrasts with the waveform distinctions of the group average QPP in these two visual conditions. This region has been linked to working memory, planning, and evaluating recency, which may be more active in the random than the systematic condition (Fincham et al., 2002; J. X. Zhang et al., 2003; Zorrilla et al., 1996). However, one should be cautious with these reverse inferences due to the significant differences between the visual stimuli used in our study and the tasks employed in previous research. Relevant to this, a significant deactivation in the default network in the PCC-precuneus was observed only in the random condition but not in the systematic condition. This significant visually-evoked deactivation was also observed in a previous study using a flickering checkerboard with a different stimulation sequence (Jorge et al., 2018). One possible explanation for the increased (decreased) engagement of the dorsolateral prefrontal region (PCC-precuneus) in the random condition observed in our study is that subjects may be less/more engaged in mind wandering during the random/systematic condition due to the focus of anticipation of the arrival of stimuli. Consequently, the default network, known to be activated during mind wandering (Godwin et al., 2017), becomes more suppressed, while the dorsolateral prefrontal region becomes more engaged during random stimulations. Among these 5 parcels, the activation of the task-negative (default) network regions and the deactivation of the task-positive network regions appears to associate with a greater amplitude of group average QPP, which remains to be investigated in the future.

### The BOLD response is dominated by the QPP waveform when visual stimulation overlaps with ongoing infraslow brain activity

Flashing checkerboards are prominent visual stimuli known to produce extensive brain activity well beyond the visual system (Gonzalez-Castillo et al., 2012; Jorge et al., 2018). In our specific experiments, across both visual conditions, nearly 70% of the BOLD stimulus responses overlap with and are overwhelmed by the waveform of ongoing primary QPPs. The BOLD response in this set is significantly greater than the 30% of trials where the stimulation does not overlap. This observation is distinct from the finding of visually stimulated BOLD response in anesthetized mice (Belloy et al., 2021), which observed the ongoing QPP only moderately affected the magnitude of subsequent stimulus BOLD (Belloy et al., 2021, Fig. 2D). This result further confirms our conjecture about the robustness of intrinsic QPP in awake humans in comparison to anesthetized mice. In other words, the dynamics of the spontaneous infraslow brain activity in the human brain that supports attention and modulates arousal is highly robust and less likely to be disrupted by environmental perturbations, though the overall dynamic waveform can be perturbed by stimulations in various ways (discussed in the 2^nd^ paragraph in this section). Moreover, the distinct patterns in the BOLD response to the different sequences of visual stimulations, demonstrated by brain parcels with significant averaged systematic-random contrast (Fig. S13), can also be captured by the distinctions of evoked ongoing QPPs between the two visual conditions. Our findings suggest that the intrinsic QPPs influenced by the flickering checkerboard may also provide a new explanation for previously reported activations and deactivations of brain regions located outside of the visual system (Jorge et al., 2018).

The widely known trial-to-trial variability in stimulated BOLD responses was popularly examined in a microscopic view in previous studies. A detailed excitation model is often described based on each stimulus and the prestimulus baseline (Chen et al., 2020; Fox et al., 2005; He, 2013; Huang et al., 2017). One influential fMRI study suggested that the observed BOLD response is a linear combination of the stimulated response and the prestimulus baseline (Fox et al., 2005). Yet, later works (Chen et al., 2020; He, 2013; Huang et al., 2017) suggest a nonadditive but inverse modulation between the stimulation and the prestimulus baseline. Specifically, a higher (lower) pre-stimulus baseline results in less (more) activation across widespread human brain regions (Huang et al., 2017) and rodent brains (Chen et al., 2020).

In these fMRI studies, even though the BOLD responses in the temporally filtered infraslow frequency range (Huang et al., 2017) or the broader low-frequency range (Fox et al., 2005; He, 2013) were studied, their underlying neurophysiological correlates remain to be investigated. The pioneering study (Chen et al., 2020) used concurrent calcium and hemodynamic imaging in the somatosensory cortical area of anesthetized rats and found a correlation between the evoked infraslow hemodynamic response and the evoked infraslow neuronal activity. However, Chen and colleagues also found that the infraslow hemodynamic power occurred before the neuronal dynamic power (Chen et al., 2020, Fig. 3ef), which implies that a significant portion of the hemodynamics may not arise from the neuronal level.

Complementing these studies with detailed activation models, our results explain this trial-to-trial variability from a macroscopic view. In particular, the varying magnitude of BOLD stimulus responses is largely controlled by the intrinsic global fluctuations of QPP – a BOLD dynamic pattern that was found to arise from the infraslow neural activity. In addition, our results provide novel insights into these non-additive activation models Chen et al., 2020; He, 2013; Huang et al., 2017). Specifically, due to the sinusoidal nature of primary QPPs (Abbas, Belloy, et al., 2019; Belloy, Shah, et al., 2018; Yousefi et al., 2018; Yousefi & Keilholz, 2021) and its window length of ~20s, the hemodynamic peak range would likely fall into a QPP phase right after the QPP phase at the prestimulus baseline, resulting in an inverse modulation between these two factors.

### Limitation and future study

The primary constraint of our study is the incomplete brain coverage caused by using a short TR and single-shot gradient EPI. This limits our ability to directly compare findings in non-visual areas to the visual system, which is most responsive to the stimuli. Additionally, while neural-BOLD adaptation to repeated visual stimuli has been well-observed in the visual cortex (Grill-Spector et al., 2006; Krekelberg et al., 2006), it is unclear how it contributes to QPPs among all brain regions. Although we did not investigate BOLD adaptation in the current study, any changes in QPPs due to neural adaptation would be reflected in the overall pattern of QPP, which is an averaged pattern across all concatenated runs. Future studies may use 7T multi-echo EPI to verify if all regions significantly activated or deactivated by visual stimuli are genuinely caused by intrinsic QPPs and if BOLD adaptation is reflected in the ongoing QPPs over time.

### Implications for BOLD fMRI

Spontaneous fluctuations in BOLD signals recorded by fMRI link to the underlying neuronal activity through complex neurovascular coupling. Yet, leveraging the multimodal imaging of BOLD and neuronal recordings, various studies reveal BOLD fluctuations directly reflecting the dynamics of neural activity in various frequency bands (Chen et al., 2020; Grooms et al., 2017; Pan et al., 2013; Raut et al., 2021; Thompson et al., 2014, 2015; X. Zhang et al., 2020). In particular, BOLD signals preserve rich information in the infralow frequency range of brain activity. This frequency range was initially dismissed as “noise”, artifact, or epi-phenomena in previous studies of circuit-level neural activity (Fox & Raichle, 2007), but has been found to have a unique neurophysiological basis closely linked to attention (Helps et al., 2010; Monto et al., 2008) and arousal (Raut et al., 2021; Sihn & Kim, 2022). More specifically, a quasi-period dynamic pattern (QPP) detected from the infraslow BOLD fluctuations was found to relate to the infraslow neuronal activity (Chen et al., 2020; Grooms et al., 2017; Pan et al., 2013; Raut et al., 2021; Thompson et al., 2014, 2015; X. Zhang et al., 2020), and also can be affected by attention (Abbas, Bassil, et al., 2019; Abbas, Belloy, et al., 2019) and arousal fluctuations (Raut et al., 2021). Thus, investigating QPP allows us to infer the dynamics of infraslow neural activity. This sheds light on understanding the interaction of environmental perturbation and evoked brain response that directly ties to the neuronal level using this noninvasive imaging technique. In addition, our results reveal that the QPP waveform is not likely to be disrupted but can still be affected by visual stimulation in various ways. This raises the possibility of developing novel non-invasive sensory stimulation procedures to perturb the dynamics of infraslow brain activity to enhance attention in humans.

## Acknowledgement

All authors thank National Science Foundation (NSF grant 1533260) for funding support. NX and SDK also thank the funding support from National Institutes of Health (NIH grant R01NS078095). DMS would like to thank the Therapeutic Cognitive Neuroscience Fund. We also thank Dr. Ying Guo for her helpful discussions.

## Supplementary Materials

The document of supplementary materials is available at https://www.biorxiv.org/content/10.1101/2022.12.06.519337v4.supplementary-material

## References

Abbas, A., Bassil, Y., & Keilholz, S. (2019). Quasi-periodic patterns of brain activity in individuals with attention-deficit/hyperactivity disorder. NeuroImage: Clinical, 21, 101653. https://doi.org/10.1016/j.nicl.2019.101653

Abbas, A., Belloy, M., Kashyap, A., Billings, J., Nezafati, M., Schumacher, E. H., & Keilholz, S. (2019). Quasi-periodic patterns contribute to functional connectivity in the brain. NeuroImage, 191, 193–204. https://doi.org/10.1016/j.neuroimage.2019.01.076

Belloy, M. E., Billings, J., Abbas, A., Kashyap, A., Pan, W.-J., Hinz, R., Vanreusel, V., Van Audekerke, J., Van der Linden, A., Keilholz, S. D., Verhoye, M., & Keliris, G. A. (2021). Resting Brain Fluctuations Are Intrinsically Coupled to Visual Response Dynamics. Cerebral Cortex, 31(3), 1511–1522. https://doi.org/10.1093/cercor/bhaa305

Belloy, M. E., Naeyaert, M., Abbas, A., Shah, D., Vanreusel, V., van Audekerke, J., Keilholz, S. D., Keliris, G. A., Van der Linden, A., & Verhoye, M. (2018). Dynamic resting state fMRI analysis in mice reveals a set of Quasi-Periodic Patterns and illustrates their relationship with the global signal. In NeuroImage (Vol. 180, pp. 463–484). Academic Press Inc. https://doi.org/10.1016/j.neuroimage.2018.01.075

Belloy, M. E., Shah, D., Abbas, A., Kashyap, A., Roßner, S., Van Der Linden, A., Keilholz, S. D. S. D., Keliris, G. A. G. A., & Verhoye, M. (2018). Quasi-Periodic Patterns of Neural Activity improve Classification of Alzheimer’s Disease in Mice. Scientific Reports, 8(1), 10024. https://doi.org/10.1038/s41598-018-28237-9

Bolt, T., Nomi, J. S., Bzdok, D., Salas, J. A., Chang, C., Thomas Yeo, B. T., Uddin, L. Q., & Keilholz, S. D. (2022). A parsimonious description of global functional brain organization in three spatiotemporal patterns. Nature Neuroscience 2022 25:8, 25(8), 1093–1103. https://doi.org/10.1038/s41593-022-01118-1

Busch, N. A., & VanRullen, R. (2010). Spontaneous EEG oscillations reveal periodic sampling of visual attention. Proceedings of the National Academy of Sciences of the United States of America, 107(37), 16048–16053. https://doi.org/10.1073/PNAS.1004801107/SUPPL_FILE/PNAS.201004801SI.PDF

Chen, W., Park, K., Pan, Y., Koretsky, A. P., & Du, C. (2020). Interactions between stimuli-evoked cortical activity and spontaneous low frequency oscillations measured with neuronal calcium. NeuroImage, 210, 116554. https://doi.org/10.1016/J.NEUROIMAGE.2020.116554

Cox, R. W. (1996). AFNI: Software for analysis and visualization of functional magnetic resonance neuroimages. Computers and Biomedical Research, 29(3), 162–173. https://doi.org/10.1006/cbmr.1996.0014

Cox, R. W., & Hyde, J. S. (1997). Software tools for analysis and visualization of fMRI data. NMR in Biomedicine, 10(4–5), 171–178. https://doi.org/10.1002/(SICI)1099-1492(199706/08)10:4/5<171::AID-NBM453>3.0.CO;2-L

Critchley, H. D., Mathias, C. J., & Dolan, R. J. (2001). Neural activity in the human brain relating to uncertainty and arousal during anticipation. Neuron, 29(2), 537–545. https://doi.org/10.1016/S0896-6273(01)00225-2

Dale, A. M., & Buckner, R. L. (1997). Selective Averaging of Rapidly Presented Individual Trials Using fMRI. Hum. Brain Mapping, 5, 329–340. https://doi.org/10.1002/(SICI)1097-0193(1997)5:5

Davis, B., Christie, J., & Rorden, C. (2009). Temporal Order Judgments Activate Temporal Parietal Junction. The Journal of Neuroscience, 29(10), 3182. https://doi.org/10.1523/JNEUROSCI.5793-08.2009

Diedenhofen, B., & Musch, J. (2015). cocor: A Comprehensive Solution for the Statistical Comparison of Correlations. PLoS ONE, 10(4). https://doi.org/10.1371/JOURNAL.PONE.0121945

Ding, J., Sperling, G., & Srinivasan, R. (2006). Attentional Modulation of SSVEP Power Depends on the Network Tagged by the Flicker Frequency. Cerebral Cortex, 16(7), 1016– 1029. https://doi.org/10.1093/CERCOR/BHJ044

Duann, J. R., Jung, T. P., Kuo, W. J., Yeh, T. C., Makeig, S., Hsieh, J. C., & Sejnowski, T. J. (2002). Single-Trial Variability in Event-Related BOLD Signals. NeuroImage, 15(4), 823– 835. https://doi.org/10.1006/NIMG.2001.1049

Engel, S. A., Glover, G. H., & Wandell, B. A. (1997). Retinotopic organization in human visual cortex and the spatial precision of functional MRI. Cerebral Cortex, 7(2), 181–192. https://doi.org/10.1093/CERCOR/7.2.181

Fincham, J. M., Carter, C. S., Van Veen, V., Stenger, V. A., & Anderson, J. R. (2002). Neural mechanisms of planning: A computational analysis using event-related fMRI. Proceedings of the National Academy of Sciences, 99(5), 3346–3351. https://doi.org/10.1073/PNAS.052703399

Fox, M. D., & Raichle, M. E. (2007). Spontaneous fluctuations in brain activity observed with functional magnetic resonance imaging. Nature Reviews Neuroscience 2007 8:9, 8(9), 700–711. https://doi.org/10.1038/nrn2201

Fox, M. D., Snyder, A. Z., Zacks, J. M., & Raichle, M. E. (2005). Coherent spontaneous activity accounts for trial-to-trial variability in human evoked brain responses. Nature Neuroscience 2005 9:1, 9(1), 23–25. https://doi.org/10.1038/nn1616

Godwin, C. A., Hunter, M. A., Bezdek, M. A., Lieberman, G., Elkin-Frankston, S., Romero, V. L., Witkiewitz, K., Clark, V. P., & Schumacher, E. H. (2017). Functional connectivity within and between intrinsic brain networks correlates with trait mind wandering. Neuropsychologia, 103, 140–153. https://doi.org/10.1016/J.NEUROPSYCHOLOGIA.2017.07.006

Gonzalez-Castillo, J., Saad, Z. S., Handwerker, D. A., Inati, S. J., Brenowitz, N., & Bandettini, P. A. (2012). Whole-brain, time-locked activation with simple tasks revealed using massive averaging and model-free analysis. Proceedings of the National Academy of Sciences of the United States of America, 109(14), 5487–5492. https://doi.org/10.1073/PNAS.1121049109/SUPPL_FILE/PNAS.201121049SI.PDF

Grill-Spector, K., Henson, R., & Martin, A. (2006). Repetition and the brain: neural models of stimulus-specific effects. Trends in Cognitive Sciences, 10(1), 14–23. https://doi.org/10.1016/J.TICS.2005.11.006

Grooms, J. K., Thompson, G. J., Pan, W. J., Billings, J., Schumacher, E. H., Epstein, C. M., & Keilholz, S. D. (2017). Infraslow Electroencephalographic and Dynamic Resting State Network Activity. Brain Connectivity, 7(5), 265. https://doi.org/10.1089/BRAIN.2017.0492

He, B. J. (2013). Spontaneous and Task-Evoked Brain Activity Negatively Interact. Journal of Neuroscience, 33(11), 4672–4682. https://doi.org/10.1523/JNEUROSCI.2922-12.2013

Helps, S. K., Broyd, S. J., James, C. J., Karl, A., Chen, W., & Sonuga-Barke, E. J. S. (2010). Altered spontaneous low frequency brain activity in Attention Deficit/Hyperactivity Disorder. Brain Research, 1322, 134–143. https://doi.org/10.1016/j.brainres.2010.01.057

Huang, Z., Zhang, J., Longtin, A., Dumont, G., Duncan, N. W., Pokorny, J., Qin, P., Dai, R., Ferri, F., Weng, X., & Northoff, G. (2017). Is There a Nonadditive Interaction Between Spontaneous and Evoked Activity? Phase-Dependence and Its Relation to the Temporal Structure of Scale-Free Brain Activity. Cerebral Cortex, 27(2), 1037–1059. https://doi.org/10.1093/CERCOR/BHV288

Jenkinson, M., Bannister, P., Brady, M., & Smith, S. (2002). Improved Optimization for the Robust and Accurate Linear Registration and Motion Correction of Brain Images. NeuroImage, 17(2), 825–841. https://doi.org/10.1006/NIMG.2002.1132

Jenkinson, M., Beckmann, C. F., Behrens, T. E. J., Woolrich, M. W., & Smith, S. M. (2012). FSL. NeuroImage, 62(2), 782–790. https://doi.org/10.1016/j.neuroimage.2011.09.015

Jones, M. R., Moynihan, H., MacKenzie, N., & Puente, J. (2002). Temporal aspects of stimulus-driven attending in dynamic arrays. Psychological Science, 13(4), 313–319. https://doi.org/10.1111/1467-9280.00458

Jorge, J., Figueiredo, P., Gruetter, R., & van der Zwaag, W. (2018). Mapping and characterization of positive and negative BOLD responses to visual stimulation in multiple brain regions at 7T. Human Brain Mapping, 39(6), 2426–2441. https://doi.org/10.1002/HBM.24012

Krekelberg, B., Boynton, G. M., & van Wezel, R. J. A. (2006). Adaptation: from single cells to BOLD signals. Trends in Neurosciences, 29(5), 250–256. https://doi.org/10.1016/J.TINS.2006.02.008

Lakatos, P., Karmos, G., Mehta, A. D., Ulbert, I., & Schroeder, C. E. (2008). Entrainment of neuronal oscillations as a mechanism of attentional selection. Science, 320(5872), 110–113. https://doi.org/10.1126/SCIENCE.1154735/SUPPL_FILE/LAKATOS.SOM.PDF

Majeed, W., Magnuson, M., Hasenkamp, W., Schwarb, H., Schumacher, E. H., Barsalou, L., & Keilholz, S. D. (2011a). Spatiotemporal dynamics of low frequency BOLD fluctuations in rats and humans. Neuroimage, 54(2), 1140–1150. https://doi.org/10.1016/j.neuroimage.2010.08.030

Majeed, W., Magnuson, M., Hasenkamp, W., Schwarb, H., Schumacher, E. H. H. E. H. H., Barsalou, L., & Keilholz, S. D. D. S. D. (2011b). Spatiotemporal dynamics of low frequency BOLD fluctuations in rats and humans. Neuroimage, 54(2), 1140–1150. https://doi.org/10.1016/j.neuroimage.2010.08.030

Majeed, W., Magnuson, M., Keilholz, S. D. S. D., & Majeed M.; Keilholz, S., W. M. (2009). Spatiotemporal dynamics of low frequency fluctuations in BOLD fMRI of the rat. J Magn Reson Imag, 30(2), 384–393. https://doi.org/10.1002/jmri.21848

Mitra, A., Kraft, A., Wright, P., Acland, B., Snyder, A. Z., Rosenthal, Z., Czerniewski, L., Bauer, A., Snyder, L., Culver, J., Lee, J. M., & Raichle, M. E. (2018). Spontaneous Infra-slow Brain Activity Has Unique Spatiotemporal Dynamics and Laminar Structure. Neuron, 98(2), 297–305.e6. https://doi.org/10.1016/j.neuron.2018.03.015

Monto, S., Palva, S., Voipio, J., & Palva, J. M. (2008). Very Slow EEG Fluctuations Predict the Dynamics of Stimulus Detection and Oscillation Amplitudes in Humans. Journal of Neuroscience, 28(33), 8268–8272. https://doi.org/10.1523/JNEUROSCI.1910-08.2008

Pan, W.-J., Thompson, G. J., Magnuson, M. E., Jaeger, D., & Keilholz, S. (2013). Infraslow LFP correlates to resting-state fMRI BOLD signals. Neuroimage, 74(0), 288–297. https://doi.org/10.1016/j.neuroimage.2013.02.035

Qiao, J., Li, X., Wang, Y., Wang, Y., Li, G., Lu, P., & Wang, S. (2022). The Infraslow Frequency Oscillatory Transcranial Direct Current Stimulation Over the Left Dorsolateral Prefrontal Cortex Enhances Sustained Attention. Frontiers in Aging Neuroscience, 14, 879006. https://doi.org/10.3389/FNAGI.2022.879006

Ramsøy, T. Z., Friis-Olivarius, M., Jacobsen, C., Jensen, S. B., & Skov, M. (2012). Effects of perceptual uncertainty on arousal and preference across different visual domains. Journal of Neuroscience, Psychology, and Economics, 5(4), 212–226. https://doi.org/10.1037/A0030198

Raut, R. V., Snyder, A. Z., Mitra, A., Yellin, D., Fujii, N., Malach, R., & Raichle, M. E. (2021). Global waves synchronize the brain’s functional systems with fluctuating arousal. Science Advances, 7(30). https://doi.org/10.1126/SCIADV.ABF2709/SUPPL_FILE/SCIADV.ABF2709_SM.PDF

Schaefer, A., Kong, R., Gordon, E. M., Laumann, T. O., Zuo, X.-N., Holmes, A. J., Eickhoff, S. B., & Yeo, B. T. T. (2018). Local-Global Parcellation of the Human Cerebral Cortex from Intrinsic Functional Connectivity MRI. Cerebral Cortex, 28(9), 3095–3114. https://doi.org/10.1093/cercor/bhx179

Schwartz, S., Vuilleumier, P., Hutton, C., Maravita, A., Dolan, R. J., & Driver, J. (2005). Attentional load and sensory competition in human vision: modulation of fMRI responses by load at fixation during task-irrelevant stimulation in the peripheral visual field. Cerebral Cortex (New York, N.Y._J: 1991), 15(6), 770–786. https://doi.org/10.1093/CERCOR/BHH178

Sedlacek, M., & Krumpholc, M. (2005). Digital measurement of phase difference - a comparative study of DSP algorithms. Metrology and Measurement Systems, *Vol.* 12(nr 4), 427–448.

Sihn, D., & Kim, S. P. (2022). Brain Infraslow Activity Correlates With Arousal Levels. Frontiers in Neuroscience, 16, 230. https://doi.org/10.3389/FNINS.2022.765585/BIBTEX

Thomas Yeo, B. T., Krienen, F. M., Sepulcre, J., Sabuncu, M. R., Lashkari, D., Hollinshead, M., Roffman, J. L., Smoller, J. W., Zöllei, L., Polimeni, J. R., Fisch, B., Liu, H., Buckner, R. L., Fischl, B., Liu, H., Buckner, R. L., Yeo, B. T., Krienen, F. M., Sepulcre, J., … Buckner, R. L. (2011). The organization of the human cerebral cortex estimated by intrinsic functional connectivity. J Neurophysiol, 106(3), 1125–1165. https://doi.org/10.1152/jn.00338.2011

Thompson, G. J., Pan, W. J., & Keilholz, S. D. (2015). Different dynamic resting state fMRI patterns are linked to different frequencies of neural activity. Journal of Neurophysiology, 114(1), 114–124. https://doi.org/10.1152/jn.00235.2015

Thompson, G. J., Pan, W. J., Magnuson, M. E., Jaeger, D., & Keilholz, S. D. (2014). Quasi-periodic patterns (QPP): Large-scale dynamics in resting state fMRI that correlate with local infraslow electrical activity. NeuroImage, 84, 1018–1031. https://doi.org/10.1016/j.neuroimage.2013.09.029

Tootell, R. B. H., Hadjikhani, N., Hall, E. K., Marrett, S., Vanduffel, W., Vaughan, J. T., & Dale, A. M. (1998). The retinotopy of visual spatial attention. Neuron, 21(6), 1409–1422. https://doi.org/10.1016/S0896-6273(00)80659-5

Urai, A. E., Braun, A., & Donner, T. H. (2017). Pupil-linked arousal is driven by decision uncertainty and alters serial choice bias. Nature Communications 2017 8:1, 8(1), 1–11. https://doi.org/10.1038/ncomms14637

Wald, L. L., & Polimeni, J. R. (2015). High-Speed, High-Resolution Acquisitions. Brain Mapping: An Encyclopedic Reference, 1, 103–116. https://doi.org/10.1016/B978-0-12-397025-1.00011-7

Wu, Q., Chang, C. F., Xi, S., Huang, I. W., Liu, Z., Juan, C. H., Wu, Y., & Fan, J. (2015). A critical role of temporoparietal junction in the integration of top-down and bottom-up attentional control. Human Brain Mapping, 36(11), 4317–4333. https://doi.org/10.1002/HBM.22919

Yousefi, B., & Keilholz, S. (2021). Propagating patterns of intrinsic activity along macroscale gradients coordinate functional connections across the whole brain. NeuroImage, 231, 117827. https://doi.org/10.1016/j.neuroimage.2021.117827

Yousefi, B., Shin, J., Schumacher, E. H., & Keilholz, S. D. (2018). Quasi-periodic patterns of intrinsic brain activity in individuals and their relationship to global signal. NeuroImage, 167, 297–308. https://doi.org/10.1016/j.neuroimage.2017.11.043

Zareian, B., Maboudi, K., Daliri, M. R., Abrishami Moghaddam, H., Treue, S., & Esghaei, M. (2020). Attention strengthens across-trial pre-stimulus phase coherence in visual cortex, enhancing stimulus processing. Scientific Reports 2020 10:1, 10(1), 1–12. https://doi.org/10.1038/s41598-020-61359-7

Zhang, J. X., Leung, H. C., & Johnson, M. K. (2003). Frontal activations associated with accessing and evaluating information in working memory: An fMRI study. NeuroImage, 20(3), 1531–1539. https://doi.org/10.1016/j.neuroimage.2003.07.016

Zhang, X., Pan, W. J., & Keilholz, S. D. (2020). The relationship between BOLD and neural activity arises from temporally sparse events. NeuroImage, 207, 116390. https://doi.org/10.1016/j.neuroimage.2019.116390

Zhang, Y., Brady, M., & Smith, S. (2001). Segmentation of brain MR images through a hidden Markov random field model and the expectation-maximization algorithm. IEEE Transactions on Medical Imaging, 20(1), 45–57. https://doi.org/10.1109/42.906424

Zhao, S., Chait, M., Dick, F., Dayan, P., Furukawa, S., & Liao, H. I. (2019). Pupil-linked phasic arousal evoked by violation but not emergence of regularity within rapid sound sequences. Nature Communications 2019 10:1, 10(1), 1–16. https://doi.org/10.1038/s41467-019-12048-1

Zhivomirov, H. (2022). *Phase Difference Measurement with Matlab*. MATLAB Central File Exchange. https://www.mathworks.com/matlabcentral/fileexchange/48025-phase-difference-measurement-with-matlab

Zorrilla, L. T., Aguirre, G. K., Zarahn, E., Cannon, T. D., & D’Esposito, M. (1996). Activation of the prefrontal cortex during judgments of recency: a functional MRI study. Neuroreport, 7(15–17), 2803–2806. https://doi.org/10.1097/00001756-199611040-00079

