## Supplementary Material for "The interaction between random and systematic visual stimulation and infraslow quasiperiodic spatiotemporal patterns of whole brain activity"

#### I. Visual stimulation sequences, motion parameters, and Schaefer-Yeo parcels coverage

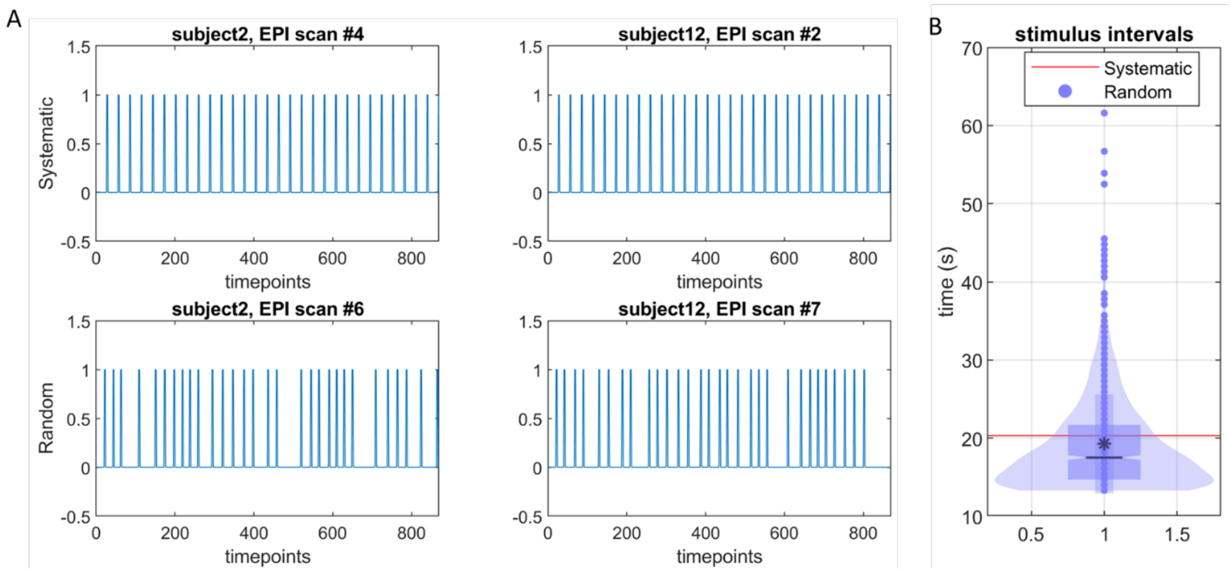


**Figure S1**: Systematic and random stimulation sequences. (A) Two examples of systematic and random stimulation sequences. (B) The violin plot of stimulus intervals for the random condition in comparison to the systematic condition for all EPI scans. Notably, the 3 systematic stimulation scans have exactly the same stimulation patterns with each stimulus arriving in every 29 TRs (20.3s, highlighted in the red horizontal line in (B)), and the 3 random stimulation scans have different stimulation patterns because each stimulus arrives at random in every 19~88 TRs (13.3s~61.6s, average 19.253s+/-6.335s). The total number of stimuli in each of the six stimulated EPI scans is always maintained to be 30.

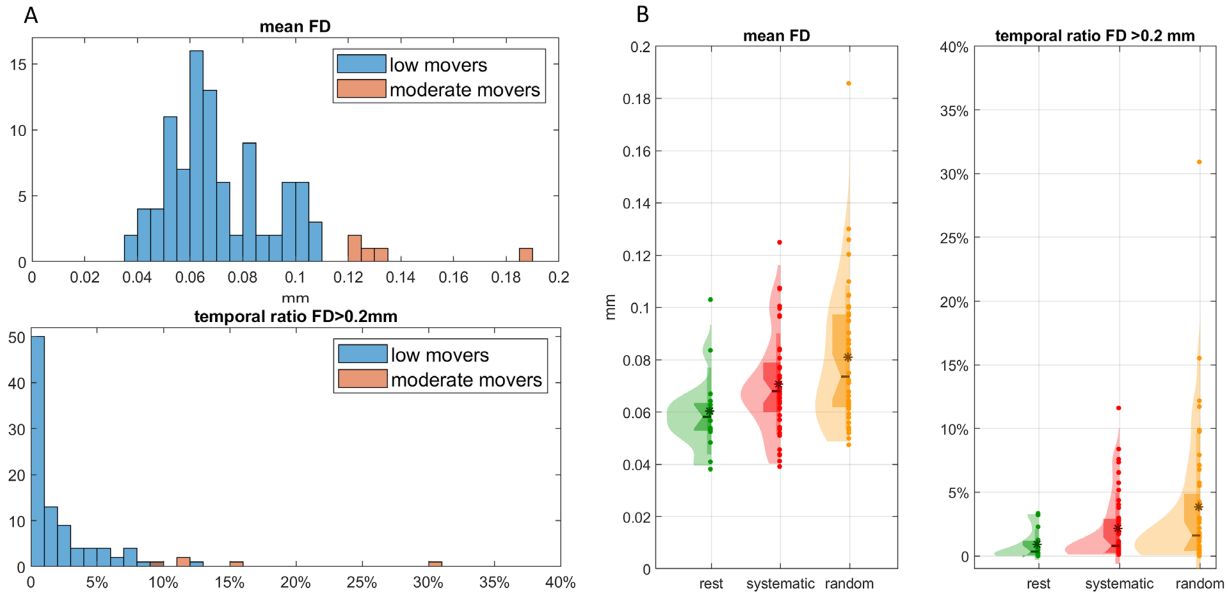


**Figure S2**: Framewise-displacement examination. (A) The histogram of mean framewise displacement (FD) (upper) and of the temporal ratio of FD>0.2mm (lower) for the total of 98 EPI scans. Among the total of 98 EPI scans (14 subjects, each has 1 resting scan, 3 systematic stimulation scans, and 3 random stimulation scans), there are 93 low moving scans, which is determined by mean framewise displacement (FD)<0.12mm (Yousefi et al., 2018), and 5 moderate moving scans (1 systematic stimulation scan and 4 random stimulation scans), which is determined by mean FD ∈ [0.12mm, 0.2mm] and the temporal ratio of FD>0.2mm spikes less than 40% (Yousefi et al., 2018). (B) The histogram of mean framewise displacement (FD) (left) and of the temporal ratio of FD>0.2mm (right) for all scans at each experimental condition. As shown, there is an increased level of motion in the systematic and then in random stimulations. Despite this increased motion level in the visual conditions, all scans only have low-moderate levels of motion, which have been found minimally impact QPPs (Yousefi et al., 2018). Hence all scans and frames were included in the group analysis of this study.

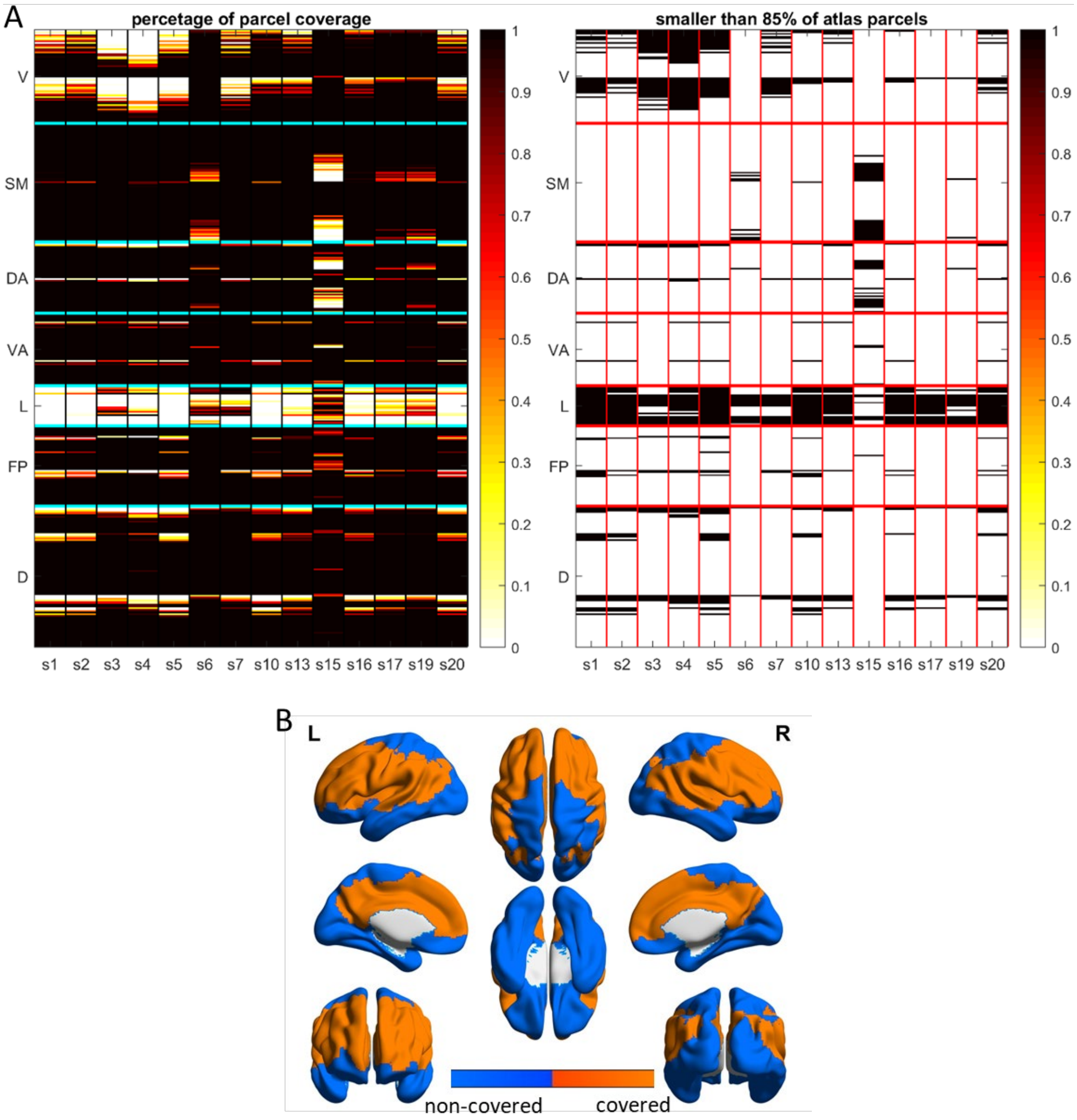


**Figure S3**: Percentage of the parcel coverage of 400 parcels from the Schaefer-Yeo atlas (A) and the brain coverage (B). In (A), parcels with more than 85% coverage in SM, DA, VA, FP, and D across all scans and all subjects (a total of 193 parcels) were included in the analysis. In (B), the brain coverage of these 193 parcels is highlighted in the brain map in orange, whereas the non-covered regions are shown in blue.

#### II. Group average QPP, reverse phase QPP, and QPP correlation timecourse

| **7 Groups** | **resting** | **s1** | **s2** | **s3** | **r1** | **r2** | **r3** |
|---|---|---|---|---|---|---|---|
| **EPI data being concatenated for each group** | 1st EPI scans of all 14 subjects | 2nd EPI scans of all 14 subjects | 4th EPI scans of all 14 subjects | 6th EPI scans of all 14 subjects | 3rd EPI scans of all 14 subjects | 5th EPI scans of all 14 subjects | 7th EPI scans of all 14 subjects |
| **group average analysis** | $QPPrest$ | All EPI scans in these three groups were used to determine $QPPsys$ | | | All EPI scans in these three groups were used to determine$QPPrand$ | | |
| independent group analysis | | $QPPsys_1$ | $QPPsys_2$ | $QPPsys_3$ | $QPPrand_1$ | $QPPrand_2$ | $QPPrand_3$ |

**Table S1:** Definition of 7 groups and their role in QPP group average analysis and independent group analysis. s1-3 are systematic 1-3 groups; r1-3 are random 1-3 groups.

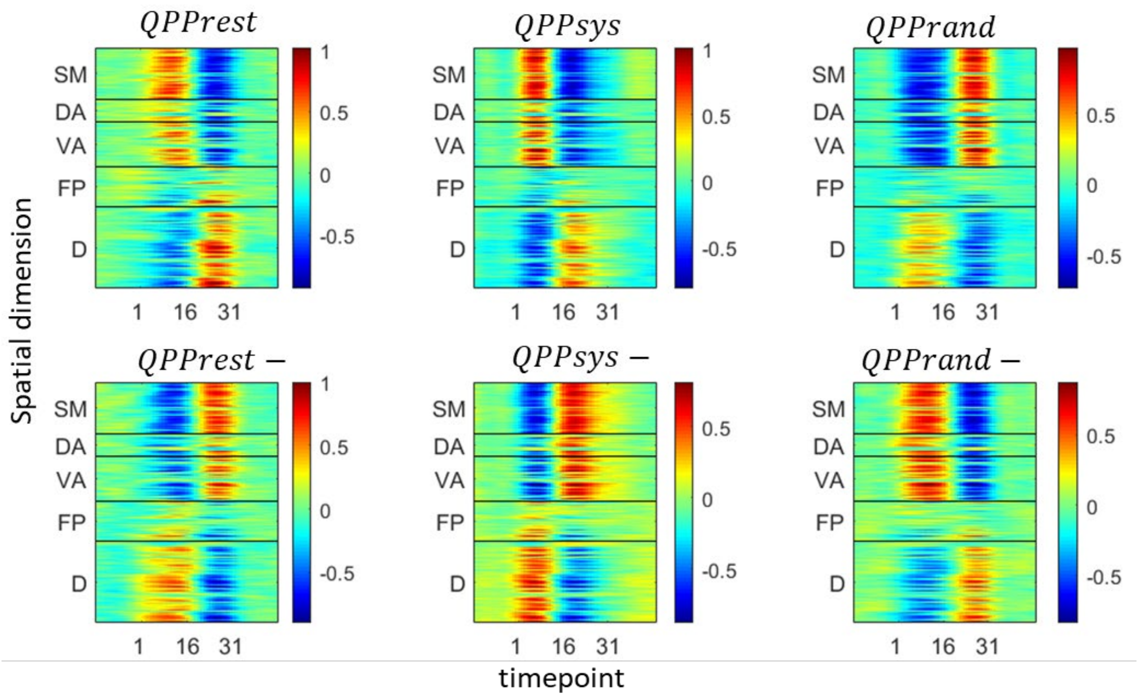


**Figure S4**: Group average QPPs and their corresponding reverse phase QPPs for resting, systematic, and random visual stimuli. The whole brain spatiotemporal patterns across five networks are shown, which include somatomotor (SM), dorsal attention (DA), ventral attention (VA), frontoparietal (FP), and default (D).

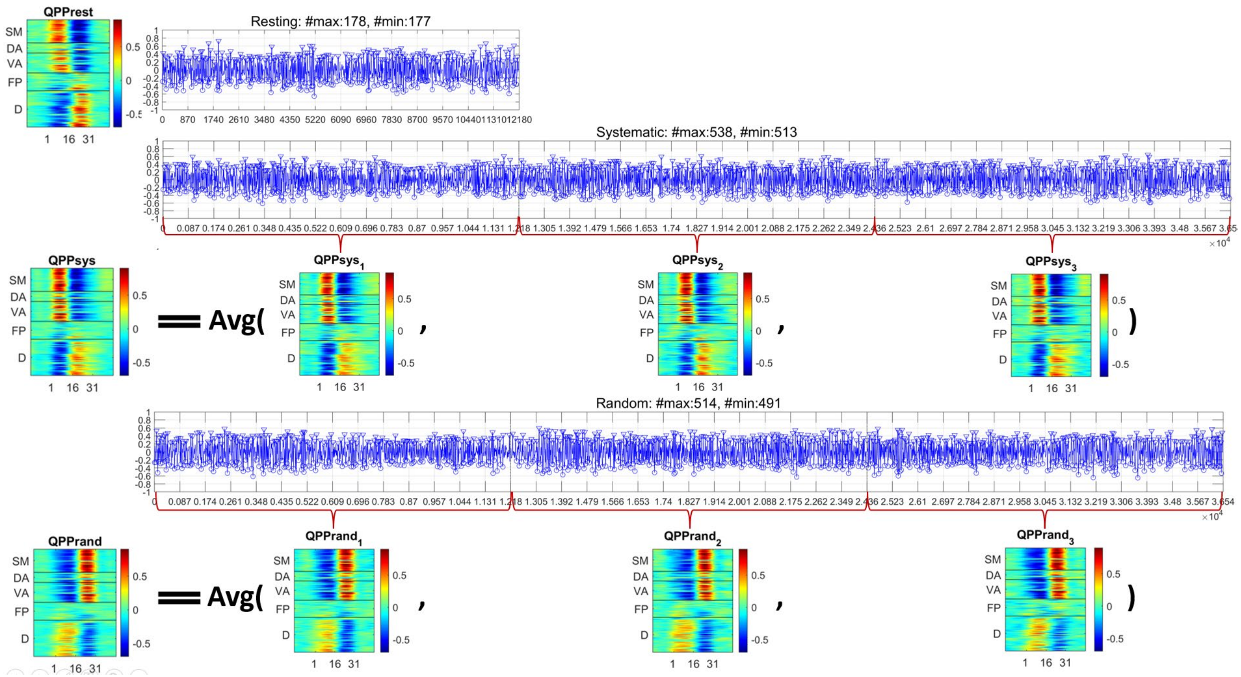


**Figure S5**: The correlation timecourse between the group average QPP template and the whole timeseries for the three experimental conditions. The primary intrinsic QPP occurred at the local maximas (labeled by triangles) whereas the phase reverse intrinsic QPP occurred at local minimas (labeled by circles) of the correlation timecourse. The QPP template for each group (sys1, sys2, sys3, rand1, rand2, and rand3) was the average of EPI segments followed by the supra-threshold peaks in the corresponding group.

| | rest | s1 | s2 | s3 | r1 | r2 | r3 |
|---|---|---|---|---|---|---|---|
| rest | 1.000 | 0.757 | 0.735 | 0.769 | 0.927 | 0.916 | 0.901 |
| s1 | 0.757 | 1.000 | 0.965 | 0.965 | NaN | NaN | NaN |
| s2 | 0.735 | 0.965 | 1.000 | 0.963 | NaN | NaN | NaN |
| s3 | 0.769 | 0.965 | 0.963 | 1.000 | NaN | NaN | NaN |
| r1 | 0.927 | NaN | NaN | NaN | 1.000 | 0.958 | 0.947 |
| r2 | 0.916 | NaN | NaN | NaN | 0.958 | 1.000 | 0.946 |
| r3 | 0.901 | NaN | NaN | NaN | 0.947 | 0.946 | 1.000 |

| | Systematic | Random |
|---|---|---|
| Average of QPP correlations between visually stimulated ($QPPsys_i$ or $QPPrand_i$) and resting ($QPPrest$ or $QPPrest -$) groups | $0.754 \pm 0.017$ | $0.918 \pm 0.012$ |
| Average of within-group QPP absolute correlations (reference) | $0.965 \pm 0.000$ | $0.953 \pm 0.007$ |

**Table S2**: Correlation of QPP between groups. QPPs for the random condition were correlated with QPPrest-. s1-3 are systematic 1-3 groups; r1-3 are random 1-3 groups. The numerical correlation values are shown in the upper table. The greater (smaller) correlation values have warmer (colder) cell colors. In the bottom table, the average correlation values for any pair of group QPPs within each visual condition are computed as the reference for the corresponding condition (2nd row), which is contrasted to the average correlation between each stimulation condition and the resting state (the 1st row). Comparing the 2nd and the 3rd column of the bottom table, the average correlation between the systematic $QPPsys_i$ and the resting QPP (r1=$0.754 \pm 0.017$, p-value<0.01) is significantly lower than the average correlation between the random $QPPrand_i$ and the resting QPP (r2=$0.918 \pm 0.012$, p-value<0.01), as the z-test statistic on the differences between these two correlation coefficients is z-score=$\frac{atanh(r1)-atanh(r2)}{\sqrt{\frac{1}{n1-3}-\frac{1}{n2-3}}}$=-32.402 (p-value<0.01). Here, the sample size for each condition is n1=n2=WL×ROI#s=31×193=5983.

| | sys | rand | LowerLimit | diff | UpperLimit | Pvalue_bfc |
|---|---|---|---|---|---|---|
| **Amplitude change** | 1 | 2 | -0.016625 | 0.058354 | 0.13333 | 0.12015 |
| **Vertical Shift** | 1 | 2 | -0.086804 | -0.061949 | -0.037093 | 4.358e-05 |
| **+"Peak-life" change** | 1 | 2 | -0.14846 | -0.047576 | 0.053312 | 0.33703 |
| **-"Peak-life" change** | 1 | 2 | -0.19516 | -0.14123 | -0.0873 | 2.3952e-05 |
| **Phase-shift** | 1 | 2 | 25.708 | 29.395 | 33.081 | 3.5315e-13 |

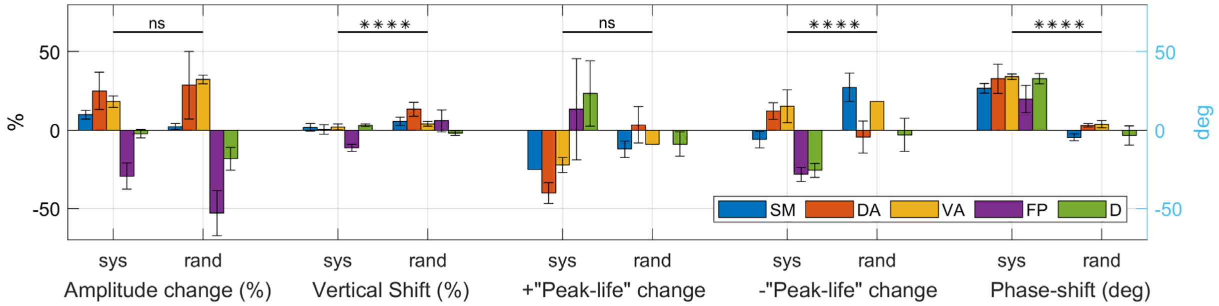


| | | SM | DA | VA | FP | D |
|---|---|---|---|---|---|---|
| Amplitude change (%) | Systematic | 9.748+/-2.811 | 24.968+/-11.937 | 18.121+/-3.718 | -29.310+/-8.225 | -2.315+/-2.679 |
| | Random | 2.322+/-2.006 | 28.613+/-21.451 | 32.326+/-2.763 | -52.926+/-14.301 | -18.300+/-7.324 |
| vertical shift (%) | Systematic | 1.781+/-2.420 | 0.410+/-3.109 | 2.012+/-1.935 | -11.371+/-2.330 | 3.089+/-0.900 |
| | Random | 5.624+/-2.702 | 13.300+/-4.358 | 3.960+/-1.570 | 5.910+/-7.077 | -1.899+/-1.459 |
| +"peak-life" change (%) | Systematic | -25.000+/-0.000 | -40.000+/-6.667 | -22.222+/-4.811 | 13.333+/-32.146 | 23.333+/-20.817 |
| | Random | -12.121+/-5.249 | 3.333+/-11.547 | -9.091+/-0.000 | NA | -8.889+/-7.698 |
| -"peak-life" change (%) | Systematic | -6.061+/-5.249 | 12.121+/-5.249 | 15.152+/-10.497 | -28.205+/-4.441 | -25.641+/-4.441 |
| | Random | 27.273+/-9.091 | -4.444+/-10.184 | 18.182+/-0.000 | NA | -3.030+/-10.497 |
| phase-shift (degree) | Systematic | 26.736+/-3.086 | 32.662+/-9.314 | 33.976+/-1.753 | 19.711+/-8.620 | 32.823+/-3.348 |
| | Random | -4.559+/-2.254 | 3.029+/-1.101 | 3.746+/-2.396 | NA | -3.281+/-6.113 |

**Table S3:** Numerical changes in QPP waveform characteristics perturbed by the visual stimulation against the resting QPP and multiple comparison test results between two visual conditions. Top: Bar plots of the average changes in three characteristics of the QPP waveform, including the amplitude changes (%), vertical shift (%), +“peak-life” change, -“peak-life” change, and phase shift (deg), as perturbed by the visual stimulation (empirical) when compared to the resting QPP (null). The significance level of the multiple comparison test between the two visual conditions is denoted above each characteristic, with 'ns' and '****' representing Bonferroni corrected p-values greater than 0.05 and less than 1e-4, respectively. The results of multiple comparison test for each characteristic are also presented. Bottom: Average changes in three characteristics of the QPP waveform. Results of the five networks including somatomotor (SM), dorsal attention (DA), ventral attention (VA), frontoparietal (FP), and default (D) are reported, which were computed from the red and black curves in Figure 3 (right). Because the negative phase for the $QPPrand$ for the frontoparietal network is almost zeros, a ‘NA’ is reported for the phase shift, the peak– change. as well as for the –“peak-life” change. Red cells indicate positive values whereas blue cells indicate negative values. The greater positive (negative) values have warmer (colder) cell colors.

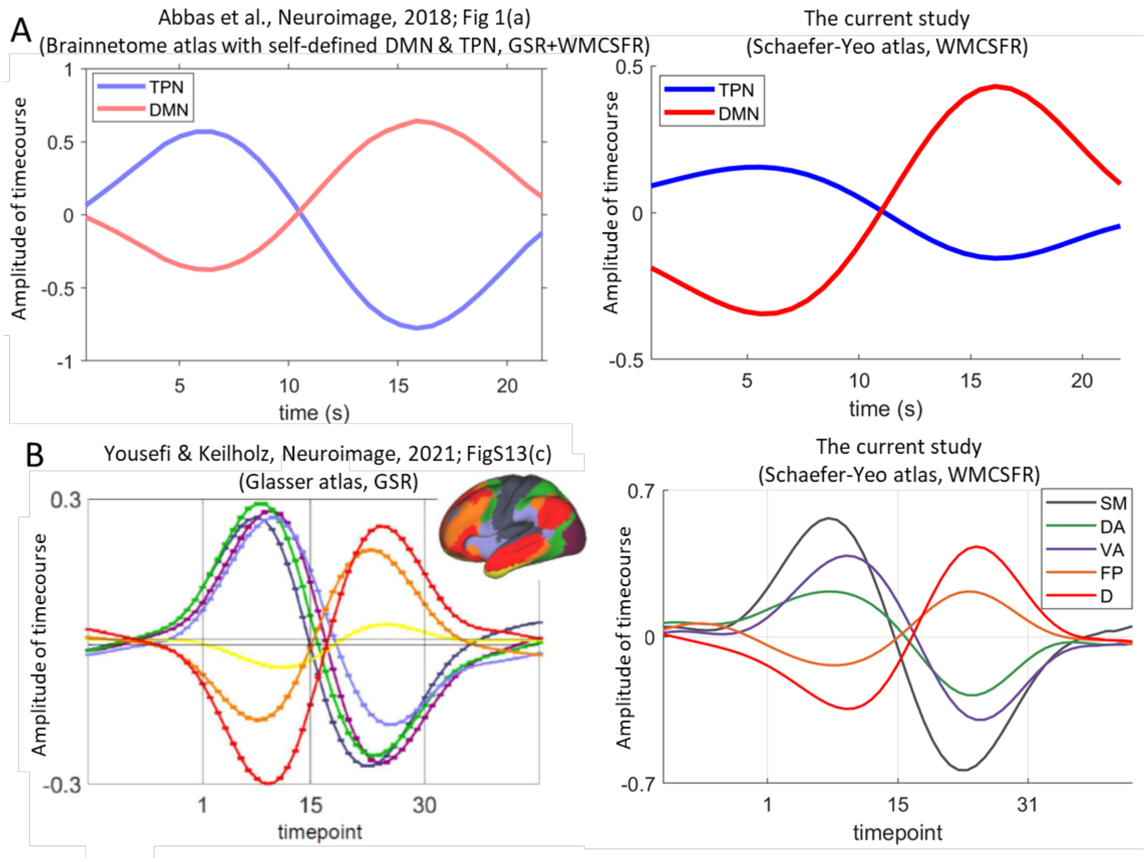


**Figure S6:** Comparison in resting-state network QPPs between different human studies. The resting QPPs of different networks of this study (right) are compared with (Abbas et al., 2019; Fig1(a)) (A, left), as well as with (Yousefi & Keilholz, 2021; FigS13 (c)) (B, left). In the above plots, the QPPs were simultaneously detected from each parcel of the whole brain, which was then averaged among parcels within the same network. In (A), the right plot captures a similar anticorrelation in QPPs between the two coarse networks (TPN and DMN) as in the left plot. In (B), the right plot captures similar correlations and anticorrelations among more fine-grained networks (Yeo et al., 2011) as depicted in the left plot. However, there are differences in the amplitude of timecourse between the left and right plots which may be due to the two distinctions in the EPI preprocessing procedures as described in the following. First, different nuisance parameter regression was applied in these studies. Specifically, (B, left) applied global signal regression (GSR), which regressed the mean signals of gray matter (GM), white matter (WM), and cerebrospinal fluid (CSF). (A, left) regressed out the mean of the entire image (also termed as one type of 'GSR') before regressing the mean of WM and CSF. In contrast, only the mean of WM and CSF signals were regressed in the current study. Second, different parcellation schemes were utilized in these three studies. For example, the (A, left) and (A, right) has slightly different definition of TPN and DMN. In (A, left), QPPs were averaged within a self-defined task-positive network (TPN) and default mode network (DMN). In particular, the TPN in (Abbas et al., 2019) mainly covers the dorsal and ventral attention areas, some visual and somatomotor areas, as well as several default and frontoparietal areas in the 7 Yeo networks (Yeo et al., 2011); the DMN in (Abbas et al., 2019) mainly covers the default and some frontoparietal areas in the 7 Yeo networks (Yeo et al., 2011). In contrast, the TPN in the current study (A, right) only encompasses the ventral and dorsal attention network plus the frontoparietal network in (Yeo et al., 2011), and the DMN (A, right) only includes the default network in (Yeo et al., 2011). On the other hand, in (B, left), the 360 Glasser parcels (Glasser et al., 2016) were clustered into the 7 Yeo networks (Yeo et al., 2011), whereas the (A, right) plot has the Yeo network based on Schaefer-Yeo parcels (Schaefer et al., 2018).

### III. Independent group QPPs

| | Sys1 | Sys2 | Sys3 | Rand1 | Rand2 | Rand3 |
|---|---|---|---|---|---|---|
| Correlation between empirical (systematic or random) and null (resting) QPPs | 0.898 | 0.650 | 0.835 | 0.929 | 0.892 | 0.772 |

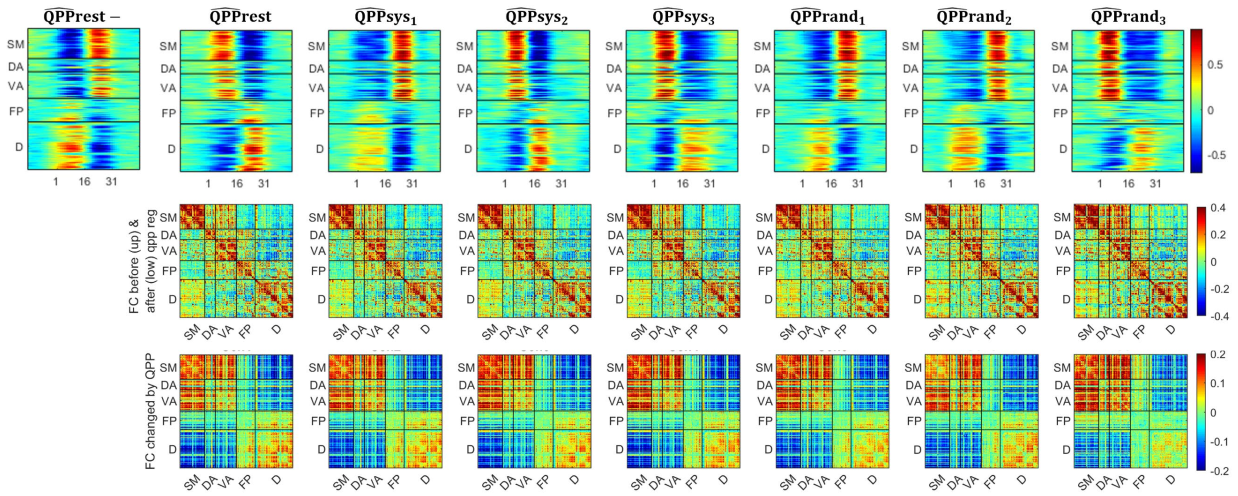


**Figure S7:** QPPs detected for each independent group (2nd row) and the correlation between visually stimulated QPP to the QPPrest or QPPrest- (top row), functional connectivity (FC) map before (upper triangular matrix of the 3rd row) and after (lower triangular matrix of the 3rd row) QPP regressions, FC map contributed by QPPs (4th row). Note that all correlation coefficients in the top row have p-values<0.01. Same to the resting populations, the primary QPP of a systematic or a random group may begin from a positive amplitude (like the sine wave) or a negative amplitude (like the -sine wave). The correlation displayed in the top row is between the stimulated QPP and the resting QPP in the same phase. The average correlation is r1=0.794+/-0.129 between systematic groups and the resting group, and it is r2=0.864+/-0.082 between random groups and the resting group. The former correlation average is significantly smaller than the latter correlation average, as the z-test statistic on the differences between these two correlation coefficients is z-score=$\frac{atanh(r1)-atanh(r2)}{\sqrt{\frac{1}{n1-3}-\frac{1}{n2-3}}}$=-12.396 (p-value<0.01). Here, the sample size for each condition is n1=n2=WL×ROI#s=31×193=5983.

| | rest | s1 | s2 | s3 | r1 | r2 | r3 |
|---|---|---|---|---|---|---|---|
| rest | 1.000 | 0.958 | 0.945 | 0.959 | 0.953 | 0.934 | 0.861 |
| s1 | 0.958 | 1.000 | 0.962 | 0.965 | 0.965 | 0.950 | 0.885 |
| s2 | 0.945 | 0.962 | 1.000 | 0.968 | 0.966 | 0.930 | 0.880 |
| s3 | 0.959 | 0.965 | 0.968 | 1.000 | 0.973 | 0.946 | 0.901 |
| r1 | 0.953 | 0.965 | 0.966 | 0.973 | 1.000 | 0.958 | 0.918 |
| r2 | 0.934 | 0.950 | 0.930 | 0.946 | 0.958 | 1.000 | 0.927 |
| r3 | 0.861 | 0.885 | 0.880 | 0.901 | 0.918 | 0.927 | 1.000 |

**Table S4**: Correlation of FC changed by QPP between groups. S1-3 are systematic 1-3 groups; r1-3 are random 1-3 groups. The greater positive (negative) values have warmer (colder) cell colors. All correlation coefficients in this table has p-value<0.01.

#### IV. Reproducibility of local QPP occurring time among independent groups

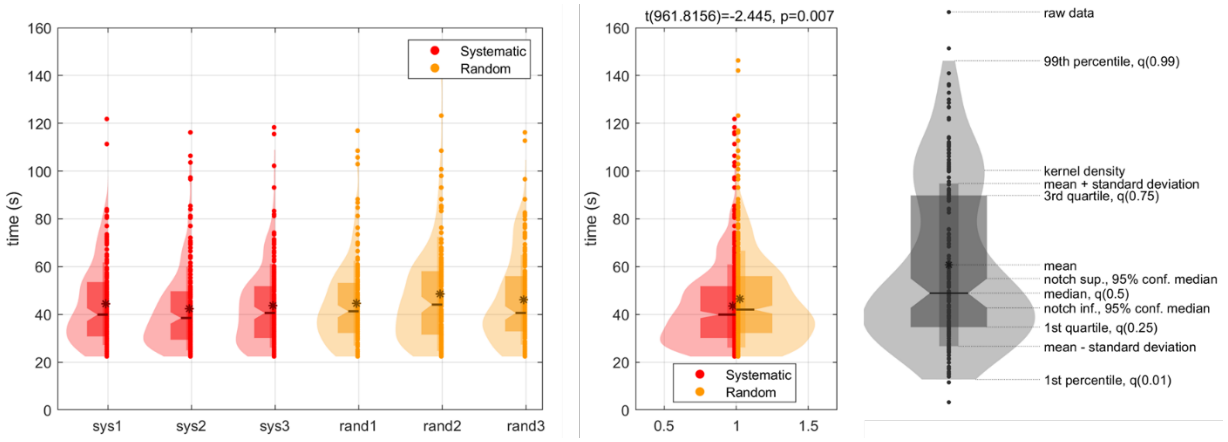


**Figure S8:** Violin plot of QPP intervals for each stimulation condition for independent groups and the manual description. QPP was independently detected for the group of subjects for every scan to increase the independency between groups. Intervals between every consecutive occurring QPP were computed and its probability density function is shown (left). The mean of QPP intervals for the 3 independent systematic groups is 44.462s, 42.461s, and 43.615s, respectively; the interval means for the 3 independent random groups are, 44.701s, 48.577s, and 46.109s, respectively. The QPP intervals were then concatenated for all 3 groups for each visual condition and then compared across two visual conditions (middle). Noted that the random intervals are always greater than the systematic ones. A one-sided t-test was performed to test the significance of such a trend. T-test statistics, including t-value(degree of freedom) and p-value, are reported. A manual description of the violin plot is also shown (right).

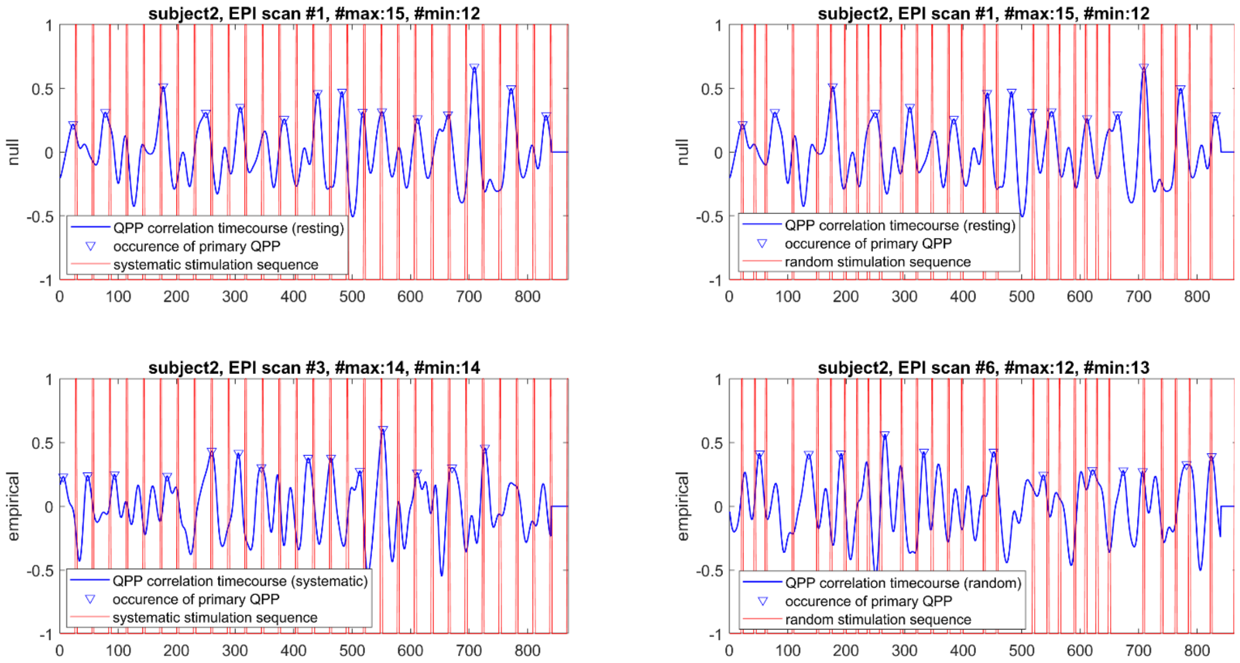


**Figure S9:** Null and empirical model of QPP delay analysis. One example of systematic and random QPP occurrence and their temporal association with the stimulation sequence is shown. The QPP correlation timecourse demonstrated in each plot is a subset extracted from the group average result.

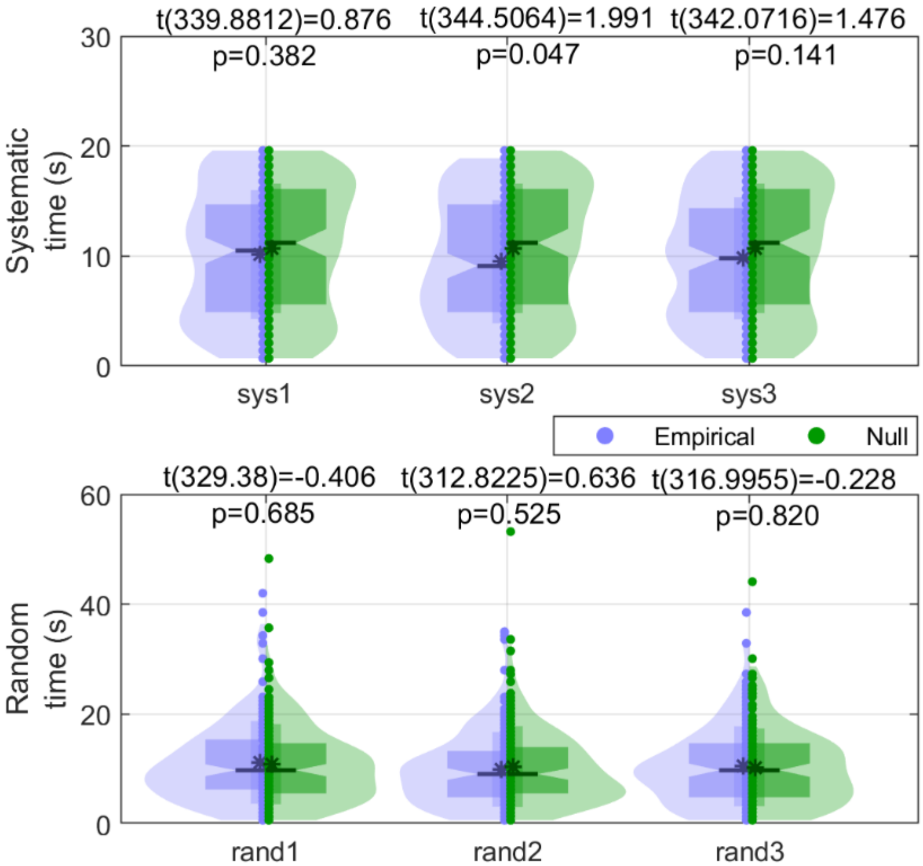


**Figure S10:** Violin plots of QPP time delay followed by visual stimuli for each independent group. QPP was independently detected for the group of subjects for every scan to increase the independency between groups. Then, for each visual condition, the QPP time delay was compared between the empirical result and the null model derived from resting data. T-tests all reject the hypothesis that the empirical mean differs from the null at the 5% significance level. For each group, t-test statistics, including t-value(degree of freedom) and p-value, are reported.

#### V. BOLD response and its dependence on QPP phases

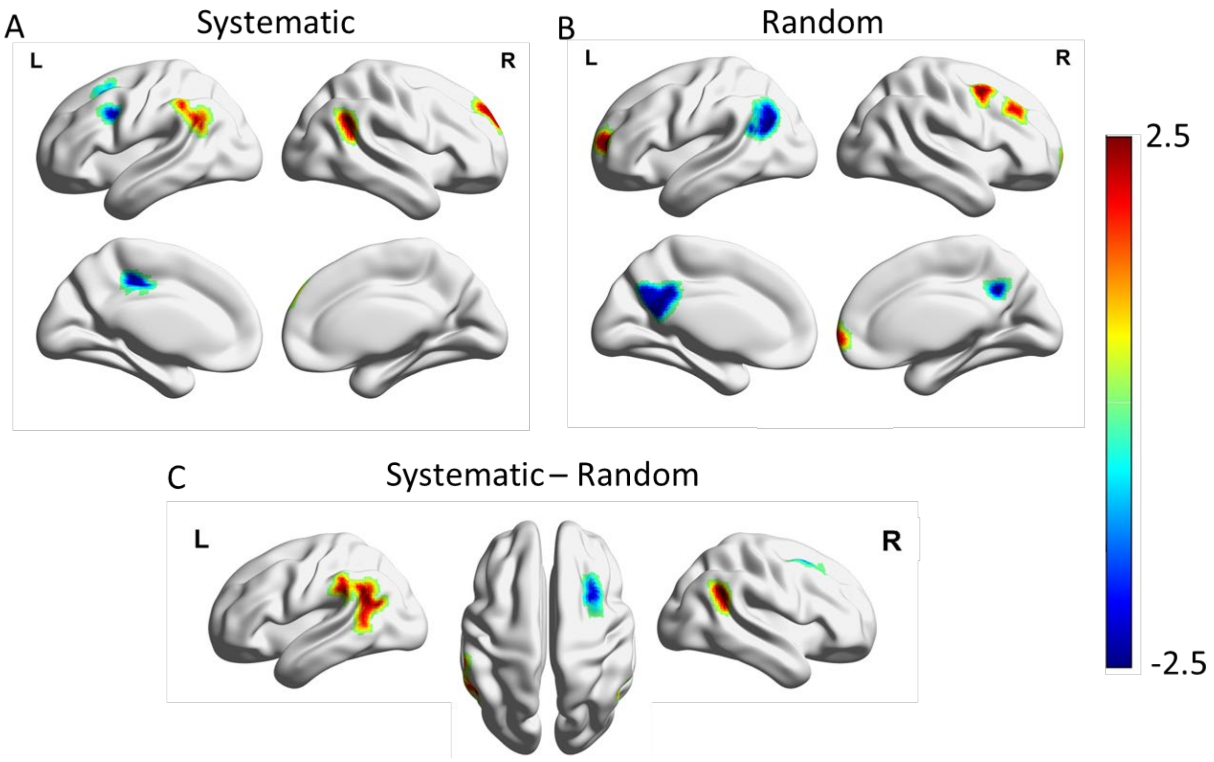


**Figure S11:** Significant BOLD peak response to systematic and random stimulations. (A) shows parcels with significant activation and deactivation (|z-score|>1.96) for the systematic condition, while (B) shows parcels with significant activation and deactivation for the random condition. (C) displays parcels with significant systematic-random contrast (|z-score|>1.96). Each plot displays the z-scores (with absolute value>1.96) of the averaged BOLD peak values. In (A) and (B), systematic visual stimulations induced significant activations in the bilateral temporoparietal junctions that spanned across the FP and D networks, as well as in the default region located in the right frontal lobe. In contrast, random visual stimulations induced significant activations in the middle frontal gyrus of the FP network as well as in the frontal pole that spanned across the FP and D networks. On the other hand, systematic stimulations resulted in significant deactivations in the frontoparietal and default regions at the middle frontal gyrus, as well as in the ventral attention region at the posterior cingulate cortex (PCC)-precentral gyrus. In contrast, random stimulations resulted in significant deactivations in the temporoparietal junctions and the PCC-precuneus region of the default network.

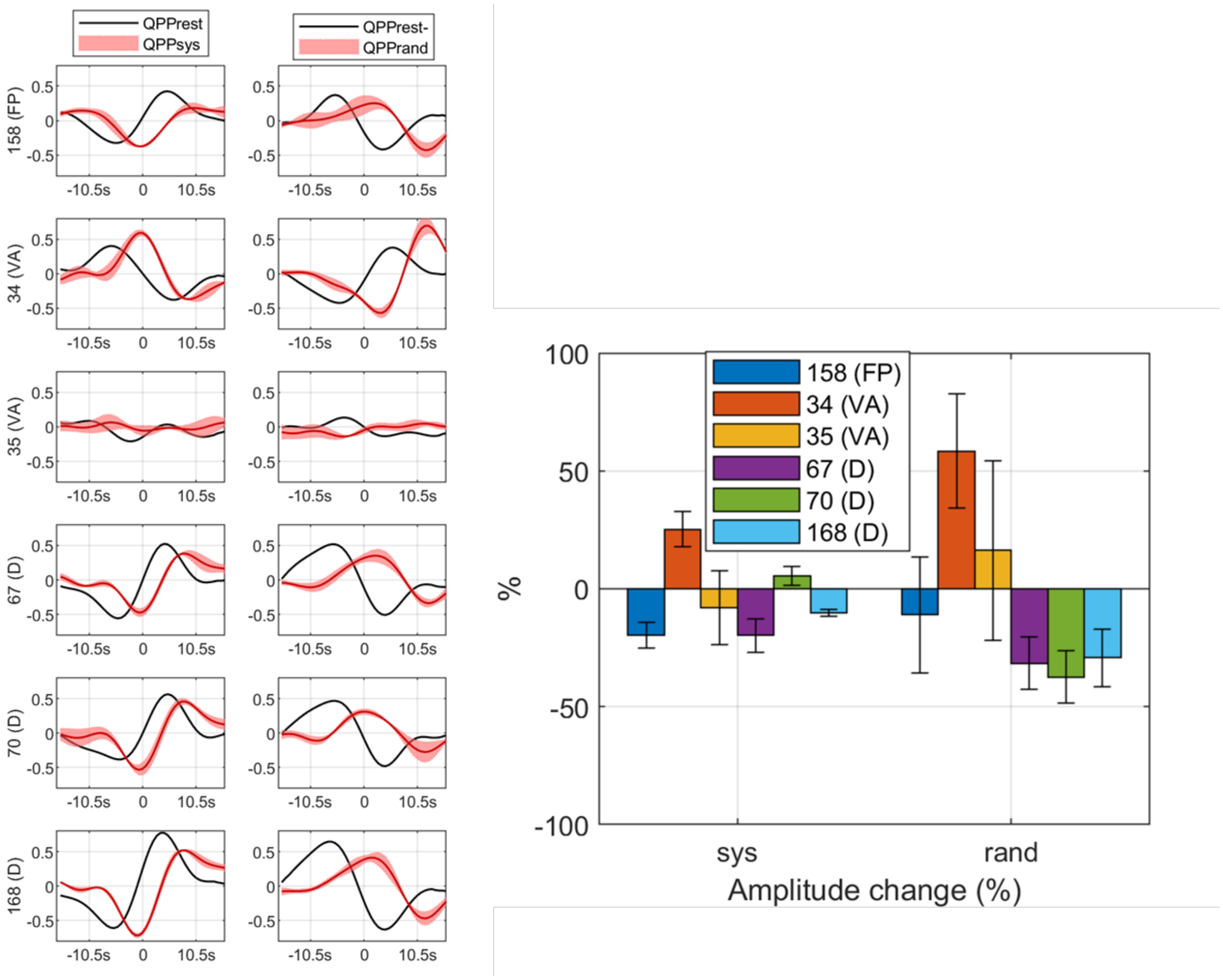


**Figure S12:** Group average QPP of resting, systematic, and random visual conditions for six parcels with significant averaged systematic-random contrast in BOLD response peaks. The y-label in the left plot and the legend in the right plot are the parcel ID and its corresponding Yeo network. The parcel ID follows (Schaefer et al., 2018)'s 400 parcels for 7 networks (webpage). These parcels demonstrate greater absolute amplitude changes by the random visual condition than by the systematic condition. In particular, the task-positive parcels (159 (FP), 34 (VA) and 35 (VA)) demonstrate much more elevated amplitude in the random than in the systematic condition; whereas the task-negative parcels (67 (D), 70 (D), and 168 (D)) demonstrate much more depressed amplitude in the random than in the systematic condition.

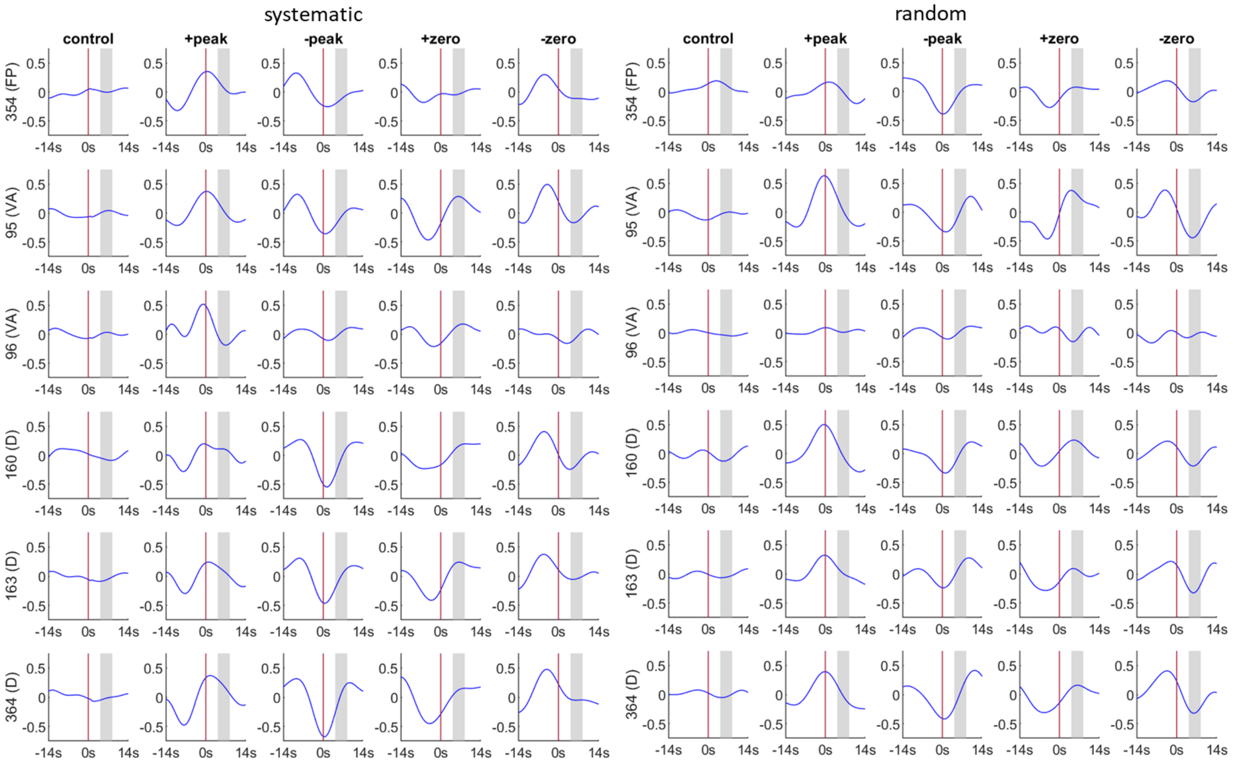


**Figure S13:** BOLD responses to systematic (upper left) and random (upper right) stimulation and their contrast (bottom left) of parcels which show significant contrasts ($p<0.01$ in t-test) between the two visual conditions. Significant 5 parcels with significant averaged systematic-random contrast in BOLD response are reported (see parcels in Fig. 5, bottom). These systematic-random contrast curves have the prestimulus baselines subtracted for each visual condition. The averaged peak value for the contrast curves (as described in Section 2.4) is reported above each plot. The parcel ID follows (Schaefer et al., 2018)'s 400 parcels for 7 networks (webpage). The control presented in the 1st column of each subfigure includes BOLD responses with no intrinsic primary QPP. The vertical axis represents the magnitude of the BOLD response whereas the horizontal axis represents the time interval before and after the stimulation occurring at 0s, which is depicted by the red vertical line. The interval between the two dotted vertical lines in each plot depicts the peak range [6TR, 12TR] of the hemodynamic response. The group average QPP for these parcels is also shown (bottom right). Note that the ranges of 4 phases were determined from the group average primary QPP of each parcel. The ongoing QPP before average may be perturbed by stimulations. For example, the systematic stimuli delay the positive peak of an ongoing QPP when they are supposed to occur at the positive peak range of the group average QPP. Such type of phase modulations is observed more in the systematic visual condition (upper left) but less in the random condition (upper right).

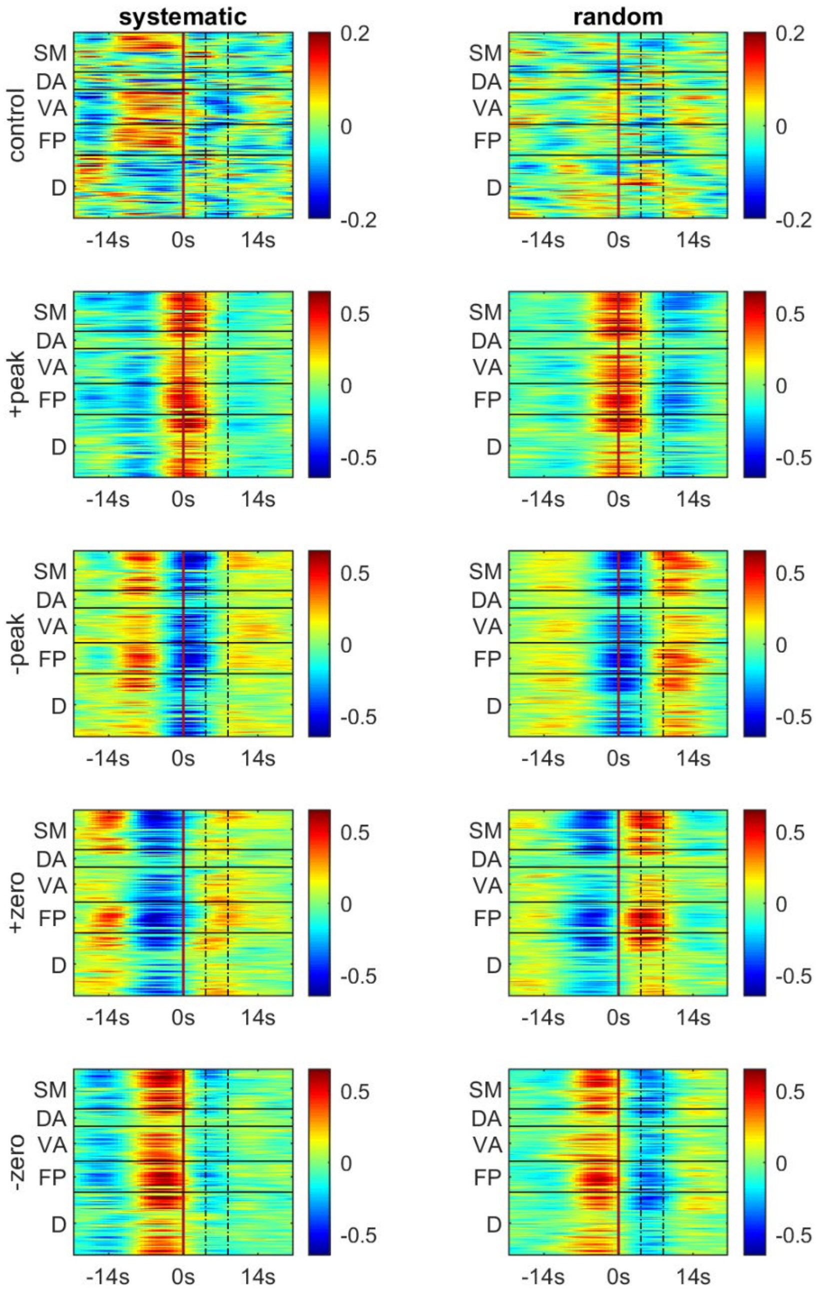


**Figure S14:** BOLD responses to systematic (left) and random (middle) stimulation of all network ROIs that are associated with four QPP phases. ROIs are stacked in rows for each network. The horizontal axis represents the time interval before and after the stimulation occurring at 0s, which is depicted by the red vertical line. The interval between the two dotted vertical lines in each plot depicts the peak range [6TR, 12TR] of the hemodynamic response.

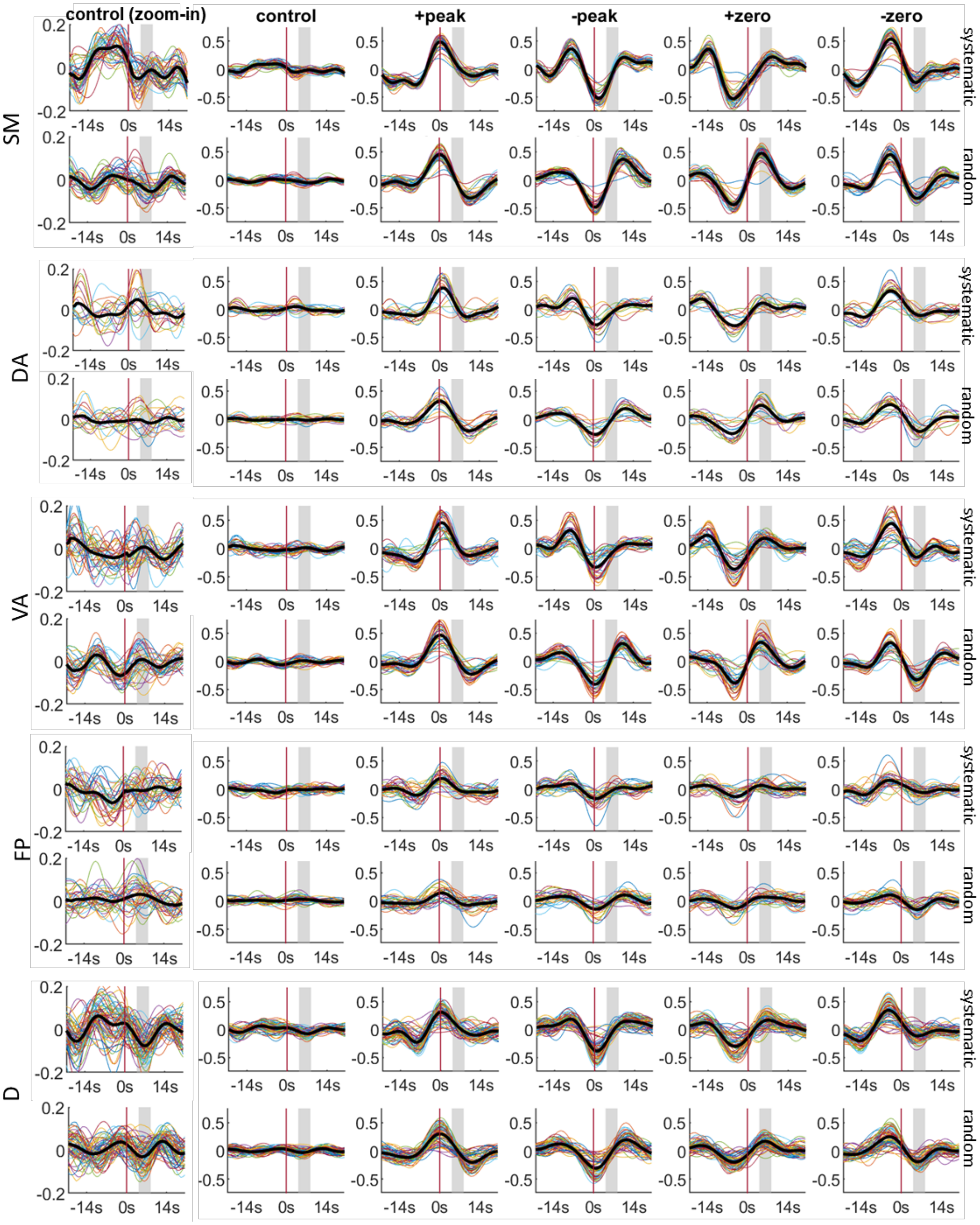


**Figure S15:** BOLD responses to systematic (upper) and random (bottom) stimulation of each network ROI that are associated with four QPP phases. ROIs are stacked in rows for each network. A zoom-in version of the control is also shown (left). In each plot, the colorful lines represent the BOLD signal of each parcel within the network, while the bold black line represents the average of all the parcels within the network. The vertical axis represents the magnitude of the BOLD response whereas the horizontal axis represents the time interval before and after the stimulation occurring at 0s, which is depicted by the red vertical line. The shaded area in each plot depicts the peak range [6TR, 12TR] of the hemodynamic response.

#### References


Abbas, A., Belloy, M., Kashyap, A., Billings, J., Nezafati, M., Schumacher, E. H., & Keilholz, S. (2019). Quasi-periodic patterns contribute to functional connectivity in the brain. *NeuroImage*, *191*, 193–204. https://doi.org/10.1016/j.neuroimage.2019.01.076

Glasser, M. F., Coalson, T. S., Robinson, E. C., Hacker, C. D., Harwell, J., Yacoub, E., Ugurbil, K., Andersson, J., Beckmann, C. F., Jenkinson, M., Smith, S. M., & Van Essen, D. C. (2016). A multi-modal parcellation of human cerebral cortex. *Nature 2016 536:7615*, *536*(7615), 171–178. https://doi.org/10.1038/nature18933

Schaefer, A., Kong, R., Gordon, E. M., Laumann, T. O., Zuo, X.-N., Holmes, A. J., Eickhoff, S. B., & Yeo, B. T. T. (2018). Local-Global Parcellation of the Human Cerebral Cortex from Intrinsic Functional Connectivity MRI. *Cerebral Cortex*, *28*(9), 3095–3114. https://doi.org/10.1093/cercor/bhx179

Yeo, B. T., Krienen, F. M., Sepulcre, J., Sabuncu, M. R., Lashkari, D., Hollinshead, M., Roffman, J. L., Smoller, J. W., Zollei, L., Polimeni, J. R., Fischl, B., Liu, H., Buckner, R. L., Thomas Yeo, B. T., Krienen, F. M., Sepulcre, J., Sabuncu, M. R., Lashkari, D., Hollinshead, M., … Buckner, R. L. (2011). The organization of the human cerebral cortex estimated by intrinsic functional connectivity. *J Neurophysiol*, *106*(3), 1125–1165. https://doi.org/10.1152/jn.00338.2011

Yousefi, B., & Keilholz, S. (2021). Propagating patterns of intrinsic activity along macroscale gradients coordinate functional connections across the whole brain. *NeuroImage*, *231*, 117827. https://doi.org/10.1016/j.neuroimage.2021.117827

Yousefi, B., Shin, J., Schumacher, E. H., & Keilholz, S. D. (2018). Quasi-periodic patterns of intrinsic brain activity in individuals and their relationship to global signal. *NeuroImage*, *167*, 297–308. https://doi.org/10.1016/j.neuroimage.2017.11.043
